# Human bronchial epithelial cell-derived extracellular vesicle therapy for pulmonary fibrosis via inhibition of TGF-β-WNT crosstalk

**DOI:** 10.1101/2020.10.22.349761

**Authors:** Tsukasa Kadota, Yu Fujita, Jun Araya, Naoaki Watanabe, Shota Fujimoto, Hironori Kawamoto, Shunsuke Minagawa, Hiromichi Hara, Takashi Ohtsuka, Yusuke Yamamoto, Kazuyoshi Kuwano, Takahiro Ochiya

**Affiliations:** Division of Respiratory Diseases, Department of Internal Medicine, The Jikei University School of Medicine, Tokyo, Japan; Division of Molecular and Cellular Medicine, National Cancer Center Research Institute, Tokyo, Japan; Division of Cellular Signaling, National Cancer Center Research Institute, Tokyo, Japan; Division of Chest Diseases, Department of Surgery, The Jikei University School of Medicine, Tokyo, Japan; Department of Molecular and Cellular Medicine, Institute of Medical Science, Tokyo Medical University, Tokyo, Japan

**Keywords:** exosome, lung epithelial cell, pulmonary fibrosis, microRNA, senescence

## Abstract

Idiopathic pulmonary fibrosis (IPF) is characterized by devastating and progressive lung parenchymal fibrosis, resulting in poor patient prognosis. An aberrant recapitulation of developmental lung gene expression, including genes for transforming growth factor (TGF)-β and WNT, has been widely implicated in the pathogenic IPF wound healing process that results from repetitive alveolar epithelial injury. Extracellular vesicles (EVs) have been shown to carry bioactive molecules and to be involved in various physiological and pathological processes. Here, we demonstrate that, by attenuating WNT signaling, human bronchial epithelial cell-derived EVs (HBEC EVs) inhibit TGF-β mediated induction of both myofibroblast differentiation and lung epithelial cellular senescence. This effect of HBEC EVs is more pronounced than that observed with mesenchymal stem cell-derived EVs. Mechanistically, the HBEC EV microRNA (miRNA) cargo is primarily responsible for attenuating both myofibroblast differentiation and cellular senescence. This attenuation occurs via inhibition of canonical and non-canonical WNT signaling pathways. Among enriched miRNA species present in HBEC EVs, miR-16, miR-26a, miR-26b, miR-141, miR-148a, and miR-200a are mechanistically involved in reducing WNT5A and WNT10B expression in LFs, and in reducing WNT3A, WNT5A, and WNT10B expression in HBECs. Mouse models utilizing intratracheal administration of EVs demonstrate efficient attenuation of bleomycin-induced lung fibrosis development accompanied by reduced expression of both β-catenin and markers of cellular senescence. These findings indicate that EVs derived from normal resident lung HBECs may possess anti-fibrotic properties. They further suggest that, via miRNA-mediated inhibition of TGF-β-WNT crosstalk, HBEC EVs administration can be a promising anti-fibrotic modality of treatment for IPF.

## Introduction

Idiopathic pulmonary fibrosis (IPF) is characterized by progressive and devastating lung parenchymal fibrosis, leading to poor patient prognosis (1). Although recently available anti-fibrotic treatment modalities, such as pirfenidone (PFD) and nintedanib, significantly reduce the decline of forced vital capacity, these drugs neither promote recovery nor prevent the progression of functional decline and the structural distortion seen in IPF (2, 3). Thus, a new therapeutic approach that substantially prevents disease progression is urgently needed. Although molecular mechanisms underlying IPF pathogenesis remain elusive, an aberrant recapitulation of developmental lung gene expression, including genes for transforming growth factor (TGF)-β and WNT, has been widely implicated in the abnormal wound healing process that follows repetitive alveolar epithelial injury (4, 5). Both TGF-β and WNT signaling are responsible for regulating myofibroblast differentiation and cellular senescence, cellular processes that have been proposed as targets with respect to IPF treatment (6, 7).

Based on the regenerative, immunomodulatory, and anti-fibrotic properties of mesenchymal stem cells (MSCs) and alveolar type II cells (ATII), injections of these cell types have gained attention as new therapeutic options for IPF. MSC administration by either intravenous or intratracheal delivery efficiently reduced inflammation and development of fibrosis in a bleomycin (BLM)-induced lung fibrosis model (8). In the context of clinical application, the safety of intravenous administration of MSCs has been demonstrated in IPF patients (9). Intratracheal administration of isolated ATII has also been shown to reduced development of BLM-induced lung fibrosis, and the safety of intratracheal administration of allogenic ATII has been evaluated in IPF patients (10, 11). Accordingly, it is likely that both MSCs and lung epithelial cells represent promising candidates for cell injection-mediated treatment of IPF. However, clinical translation of cell injection therapies has been fraught with technical challenges in terms of scalability, reproducibility, and safety. Furthermore, the precise mechanisms of action underlying cell injection therapies remain uncertain. Due to the prominent role of the secretome in many biological processes, secretome-based therapies, including the use of extracellular vesicles (EVs), have been proposed as an efficacious alternative to cell injection strategies (12).

EVs are double-layered lipid membrane vesicles that are categorized as exosomes, microvesicles, and apoptotic bodies, based on their size, biogenesis and secretory mechanisms (13). EVs are important mediators of intercellular communication via the transfer of their bioactive cargo molecules such as microRNAs (miRNAs) and proteins (14). MSC-derived EVs (MSC EVs) are recognized as being at least partly responsible for MSC-mediated regeneration and immunomodulation (15). Furthermore, growing evidence suggests that MSC EVs represent a promising cell-free therapeutic approach for a wide array of disorders, including not only inflammatory lung disorders, but also lung fibrosis (12, 16–18). The anti-fibrotic properties of ATII cell therapy indicate the potential effectiveness of ATII-derived EVs (ATII EVs) for IPF treatment. However, there is no appropriate *in vitro* ATII culture system capable of preparing sufficient quantities of ATII EVs for injection therapy. Intriguingly, a recent paper demonstrates the anti-fibrotic property of lung spheroid-derived EVs generated from healthy lung samples (19, 20). Our recent paper shows that the miRNA cargo of EVs derived from cigarette smoke (CS)-exposed human bronchial epithelial cells (HBEC EVs) can induce myofibroblast differentiation associated with airway remodeling. This suggests the potential regulatory role of airway epithelial cell (AEC)-derived EVs (AEC EVs) in development of lung fibrosis (21). Because of the result with damaged HBECs, we have hypothesized that healthy AEC EVs may have the opposite effect; i.e. they may possess anti-fibrotic properties that are valuable for IPF treatment.

In the present study, anti-fibrotic mechanisms mediated by AEC EVs, including human small airway epithelial cells (HSAECs) and HBECs, are evaluated using *in vitro* models of TGF-β-induced myofibroblast differentiation and cell senescence. The details of anti-fibrotic mechanisms mediated by HBEC EVs are analyzed in the context of their miRNA cargo. The anti-fibrotic properties of HBEC EVs are further examined *in vivo* via intratracheal administration of these vesicles in mouse models of BLM-induced lung fibrosis.

## Results

### HBEC-derived EVs attenuate TGF-β-induced myofibroblast differentiation and lung epithelial cell senescence

HBECs and HSAECs were isolated from bronchus and lung parenchyma, respectively (Fig. 1A, Table S1). EVs derived from these cells were isolated by conventional ultracentrifugation of conditioned medium collected from primary HBECs, HSAECs and from immortalized HBECs (BEAS-2B cells). EVs were characterized by western blotting to identify exosome markers and by nanoparticle tracking analysis (NTA) (Figs. S1A, B) (21). These EVs fulfilled the minimal experimental criteria for identification as exosomes, as described in the position statement from the International Society for Extracellular Vesicles (22). To characterize the anti-fibrotic properties of these EVs in terms of IPF treatment(23), we uses lung fibroblasts (LFs) to examine EV effects on TGF-β-induced myofibroblast differentiation (Fig. 1B). TGF-β-induced myofibroblast differentiation, as judged by increased expression of type I collagen and αSMA, was clearly suppressed by BEAS-2B EVs, HBEC EVs, and HSAEC EVs (Fig. 1B). Fig. 1C uses immunofluorescence labeling of type I collagen and αSMA to illustrate the inhibitory effect of AEC EVs on TGF-β-induced myofibroblast differentiation. In comparison to HSAECs, significantly higher numbers of EVs were secreted from HBECs and BEAS-2B cells (Figs. S1C-E). In addition, HBEC EVs exhibited the strongest dose-dependent inhibitory activity against myofibroblast differentiation, consistent with possible anti-fibrotic properties of HBEC EVs (Figs. 1B, 1E, Fig. S1G). To clarify the specific activity of AEC EVs in suppressing myofibroblast differentiation, we also examined the anti-fibrotic properties of EVs derived from other cell types, including MSCs (Fig. 1D, Fig. S1F). EVs derived from LFs, Human umbilical vein endothelial Cells (HUVEC), a monocyte cell line (THP-1), and lung cancer cell lines (A549, PC9, PC14) did not attenuate TGF-β-induced type I collagen and αSMA expression. Intriguingly, AEC EVs, especially HBEC EVs, more efficiently suppressed TGF-β-induced myofibroblast differentiation than bone marrow MSC (BM-MSC)-derived EVs. (Fig. 1D, Fig. S1F). The strength of the inhibitory effect of HBEC EVs on myofibroblast differentiation was compared with those of PFD and nintedanib (Fig. 1E, Fig. S1G). The approximate maximum drug concentrations (Cmax) in clinical use is 10 μg/ml for PFD and 60 nM for nintedanib. Compared to dosed of 10 μg/ml PFD and 100nM nintedanib, HBEC EVs exhibited more effective suppression of TGF-β-induced myofibroblast differentiation in LFs (Fig. 1E, Fig. S1G).

**Figure 1.**
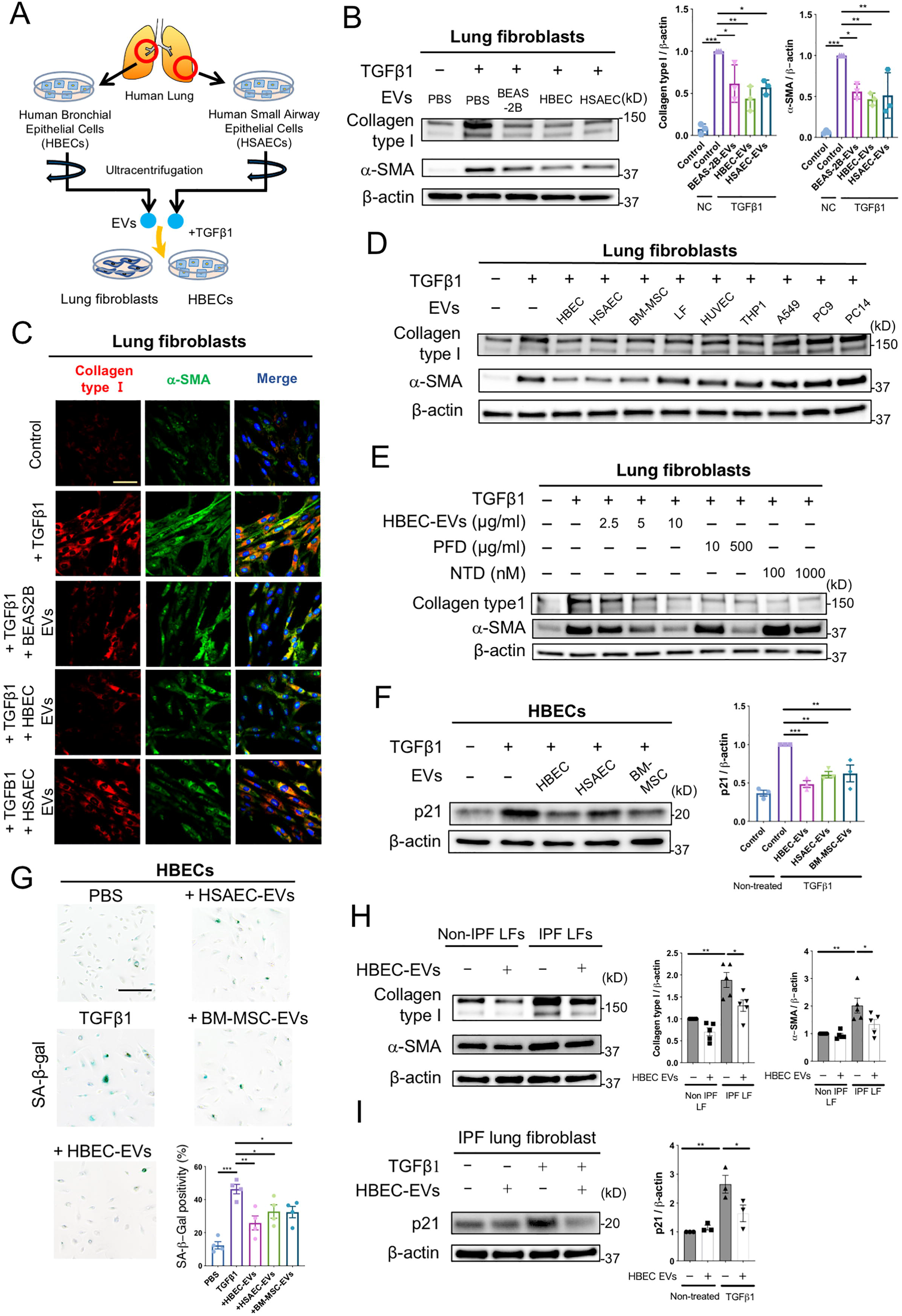
Lung epithelial cell-derived EVs attenuate TGF-β-induced myofibroblast differentiation and lung epithelial cell senescence. **A** Schematic protocol for the administration of airway epithelial cell-derived EVs, including human bronchial epithelial cell (HBEC) EVs and human small airway epithelial cell (HSAEC) EVs, to lung fibroblasts (LFs) or HBECs. **B** Representative immunoblot and quantitative analysis showing the amount of type I collagen, α-smooth muscle actin (SMA), and β-actin in LFs treated for 24 h with PBS, BEAS-2B EVs (10 μg/ml), HBEC EVs (10 μg/ml) or HSAEC EVs (10 μg/ml) in the presence or absence of TGF-β1 (2 ng/ml). Protein samples were collected from treated cells 72 h after initiation of stimulation. *p<0.05, **p<0.005, ***p<0.001. **C** Representative images of anti-α-SMA, anti-type I collagen and Hoechst 33342 staining in LFs incubated for 24 h with PBS, BEAS-2B EVs (10 μg/ml), HBEC EVs (10 μg/ml) or HSAEC EVs (10 μg/ml) in the presence or absence of TGF-β1 (2 ng/ml). Staining was performed 72 h after beginning stimulation. Bar: 20 μm. **D** Representative immunoblot showing the amount of type I collagen, α-SMA, and β-actin in LFs treated for 24 h with PBS, HBEC EVs, HSAEC EVs, BM-MSC EVs, LF EVs, HUVEC EVs, THP1 EVs, A549 EVs, PC9 EVs, or PC14 EVs in the presence or absence of TGF-β1 (2 ng/ml). EVs were added to the medium at a concentration of 2 × 10^9^ particles per ml. Protein samples were collected after 72 h of stimulation. **E** Representative immunoblot showing the amount of type I collagen, α-SMA, and β-actin in LFs treated for 24 h with PFD (10 or 500 μg/ml) or nintedanib (NTD) (100 or 1000 nM) in the presence or absence of TGF-β1 (2 ng/ml). Protein samples were collected 72 h after stimulation. **F** Representative immunoblot showing the amount of p21 and β-actin in HBECs treated for 48 h with PBS, HBEC EVs, HSAEC EVs and BM-MSC EVs in the presence or absence of TGF-β1 (2 ng/ml). EVs were added to the medium at a concentration of 2 × 10^9^ particles per ml. Protein samples were collected 96 h after beginning stimulation. **p<0.005, ***p<0.001. **G** Representative images of SA-β-gal staining in HBECs. HBECs were treated for 48 h with PBS, HBEC EVs, HSAEC EVs and BM-MSC EVs in the presence or absence of TGF-β1 (2 ng/ml). EVs were added to the medium at a concentration of 2×10^9^ particles per ml. Staining was performed 96 h after the start of treatment. *p<0.05, **p<0.005, ***p<0.001. Scale bar, 200 μm. **H** Representative immunoblot showing the amount of type I collagen, α-SMA, and β-actin in IPF LFs treated for 48 h with PBS or HBEC EVs (10 μg/ml). *p<0.05, **p<0.005. **I** Representative immunoblot showing the amount of p21 and β-actin in IPF LFs treated for 48 h with PBS or HBEC EVs (10 μg/ml) in the presence or absence of TGF-β1 (2 ng/ml). Protein samples were collected 96 h after beginning stimulation. *p<0.05, **p<0.005.

To further explore the possible anti-fibrotic nature of HBEC EVs, the ability of these EVs to protect against cellular senescence was examined using lung epithelial cells. It has been reported that senescent metaplastic epithelial cells covering fibroblastic foci (FF) originated mainly from bronchioles in IPF lungs (24), and bronchiolization in distorted parenchyma is a characteristic pathologic finding in IPF lungs. In the context of IPF pathogenesis, TGF-β is known to induce lung epithelial cell senescence in a p21 dependent manner (25). We therefore selected HBECs to examine the effect of EVs on cellular senescence. Interestingly, treatment with HBEC EVs clearly inhibited TGF-β-mediated induction of p21 expression (Fig. 1F, Fig. 1H). In addition, TGF-β-induced expression of senescence-associated-β-galactosidase (SA-β-gal) staining was also suppressed by HBEC EVs (Fig. 1G). Compared to HSAEC EVs and MSC EVs, HBEC EVs more efficiently suppressed TGF-β-induced cellular senescence (Figs. 1F, G). Recent studies including our own indicate that IPF lung-derived LFs (IPF LFs), in contrast to non-IPF lung-derived LFs (Non-IPF LFs), are phenotypically characterized not only by myofibroblast differentiation but also by cellular senescence associated with increased p21 and p16 expression (26, 27). Accordingly, we evaluated the effect of HBEC EVs on myofibroblast differentiation and cellular senescence in IPF LFs (Table S2). Intriguingly, HBEC EVs significantly suppressed myofibroblast differentiation in IPF LFs (Fig. 1H) and efficiently inhibited TGF-β-induced cellular senescence in IPF LFs (Fig. 1I). Together, these data suggest that the anti-fibrotic properties of HBEC EVs can be attributed to suppression of both myofibroblast differentiation and cellular senescence caused by TGF-β.

### HBEC EVs suppress TGF-β-induced activation of both canonical and non-canonical WNT signaling pathways in LFs

To clarify the molecular mechanisms underlying HBEC EV- and BEAS-2B EV-mediated suppression of TGF-β-induced myofibroblast differentiation, we looked for alterations in TGF-β-evoked signaling pathways in response to EV treatment of LFs. Our investigation focused on pathways related to SMAD2/3, PI3K/AKT, MAPK, WNT signaling, and NOX4 signaling. Consistent with previous findings, TGF-β treatment of LFs clearly activated SMAD2/3, PI3K/AKT, MAPK, and WNT signaling, and induced NOX4 expression (Fig. S2A). Among these potential targets, only canonical WNT signaling, as judged by expression of β-catenin, was significantly suppressed by both HBEC EVs and BEAS-2B EVs (Fig. 2A, Figs. S2A, B). No suppression of SMAD2/3 signaling was detected, and the MAPK, AKT, and NOX4 were slightly activated by HBEC EVs (Fig. S2A). To confirm the HBEC EV-mediated regulation of canonical WNT signaling, nuclear translocation of β-catenin was examined by means of nuclear/cytoplasmic fractionation. TGF-β treatment of LFs clearly upregulated β-catenin protein levels in both the cytoplasmic and nuclear fractions. HBEC EV treatment significantly suppressed β-catenin levels in both of these fractions (Fig. 2B).

**Figure 2.**
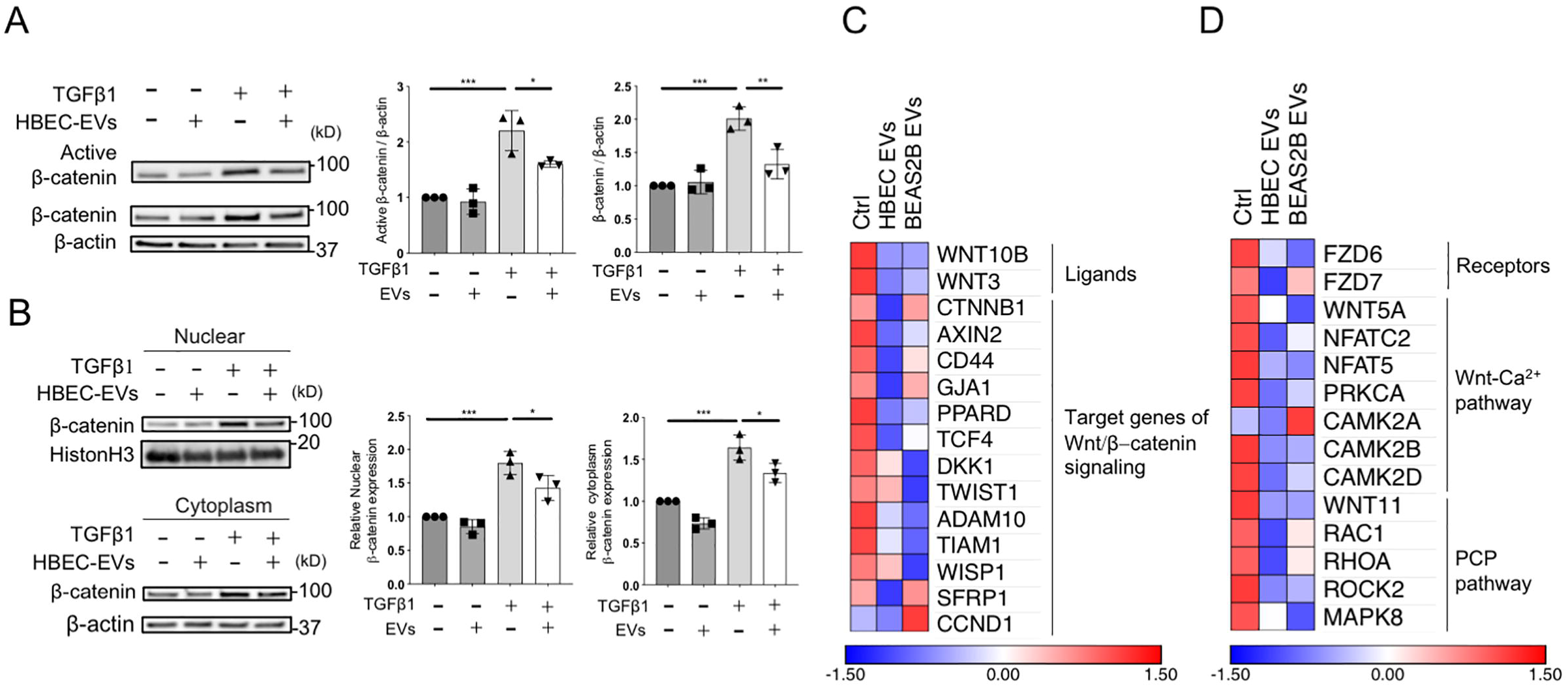
HBEC EVs attenuate myofibroblast differentiation via canonical and non-canonical WNT signaling pathways. **A** Representative immunoblot and quantitative analysis showing the amount of active β-catenin, β-catenin and β-actin in LFs treated for 48 h with PBS or HBEC EVs (10 μg/ml) in the presence or absence of TGF-β1 (2 ng/ml). *p<0.05, **p<0.005, ***p<0.001. **B** Representative immunoblot and quantitative analysis showing β-catenin and histone H3 in LFs treated for 48 h with PBS or HBEC EVs (10 μg/ml) in the presence or absence of TGF-β1 (2 ng/ml). *p<0.05, ***p<0.001. **C** Heat map of RNA-seq results showing differences in expression of Wnt/β-catenin signaling elements in LFs treated for 48 h with PBS, HBEC EVs (10 μg/ml) or BEAS-2B EVs (10 μg/ml). **D** A heat map of RNA-seq results showing differences in expression of Wnt-Ca^2+^ pathway and planar cell polarity pathway elements in LFs treated for 48 h with PBS, HBEC EVs (10 μg/ml) or BEAS-2B EVs (10 μg/ml).

To further identify target molecules in the suppression of TGF-β-induced myofibroblast differentiation by HBEC EVs and BEAS-2B EVs, we performed RNA sequencing (RNA-seq) analysis of LFs treated with HBEC EVs or BEAS-2B EVs. Consistent with observed protein levels, transcription of message for fibrogenic components, including ACTA2, COL1A1 and COL1A2, was downregulated by treatment with HBEC EVs and BEAS-2B EVs (Fig. S2C). Gene set enrichment analyses (GSEA) of the RNA-seq data revealed that transcription of genes associated with the TGF-β pathway, extra matrix, extracellular matrix structural constituent, and collagen fibril organization was significantly downregulated by HBEC EVs and BEAS-2B EVs compared to control treated LFs (Fig. S2D, E). In line with evaluation of TGF-β associated signaling by western blotting (Figs. 2A, B), RNA-seq demonstrated reduced transcription of ligands and target genes associated with canonical WNT/β-catenin signaling in LFs treated with HBEC EVs or BEAS-2B EVs (Figs. 2C, D). In addition, these EVs also suppressed transcription of essential components of the non-canonical WNT-Ca^2+^ and planar cell polarity (PCP) pathways. These findings suggest the involvement of both canonical and non-canonical WNT signaling pathways in the mechanisms responsible for HBEC EVs and BEAS-2B EVs mediated suppression of TGF-β-induced myofibroblast differentiation.

### MiRNA cargo in HBEC EVs is responsible for attenuation of TGF-β-induced myofibroblast differentiation

The biomolecular cargo in HBEC EVs was examined to further clarify the molecular mechanisms by which HBEC EVs attenuate TGF-β induction of both canonical and non-canonical WNT signaling pathways. Since EVs contain a variety of biomolecules, including proteins and miRNAs, we first performed proteomic analysis of HBEC EVs by means of mass spectrometry (MS) (Table S3). The purity of the EV preparation was confirmed by assessment of EV markers, according to a MISEV2018 position paper (Fig. S3A) (22). To understand the biological function of proteins contained in HBEC EVs, we used DAVID 6.8 to perform gene ontology (GO) enrichment analysis and Kyoto Encyclopedia of Genes and Genomes (KEGG) pathway analysis of co-expressed proteins in two different HBEC EV preparations (Fig. S3B). Reflecting the characteristics of EVs, most of the proteins identified were exosomal, cytosolic, and membranous in nature (Fig. S3C). Beyond these elements, molecular function analysis suggested that HBEC EV cargo was significantly enriched for proteins responsible for metabolic and catabolic processes (Figs. S3D, E). However, no apparent enrichment for co-expressed proteins relevant to TGF-β and WNT signaling was detected in our GO and KEGG pathway analysis (Figs. S3D, E). Interestingly, greater HBEC EV suppression of myofibroblast differentiation was observed after 72 h incubation than after 24h or 48 h. This supports the notion that the effect of EVs might be mediated via not receptor-ligand interaction, but miRNA transfer (Fig. S3F). Hence, we performed miRNA RNA-seq analysis with these EVs, focusing on the 30 most abundant miRNAs present in HBEC EVs (Figs. 3A, B). We based our analysis on miRNA microarray datasets GSE32358 in the Gene Expression Omnibus (GEO) database, including 106 IPF lungs and 50 healthy lungs. Our study revealed that 25 of the 30 miRNAs highly-expressed in HBEC EVs were downregulated in IPF lung samples, whereas 5 miRNAs were upregulated (Fig. 3C). To determine the functional properties of the top 30 miRNAs present in HBEC EVs, we used DIANA-mirPath analysis to perform KEGG pathway and GO analyses. KEGG pathway analysis indicated that the 30 miRNAs were involved in regulating TGF-β and WNT signaling (Fig. 3D). GO analysis revealed significant enrichment for miRNAs involved in biological processes such as extracellular matrix organization, aging, collagen fibril organization, collagen catabolic processes, cell death, and cell cycle arrest (Fig. 3E). Taken together, these findings suggest that the 30 miRNAs present in HBEC EVs negatively regulate TGF-β signaling, with concomitant effect on WNT-related pathways. The overall effect of this negative regulation could be responsible for increases in fibrosis and cellular senescence.

**Figure 3.**
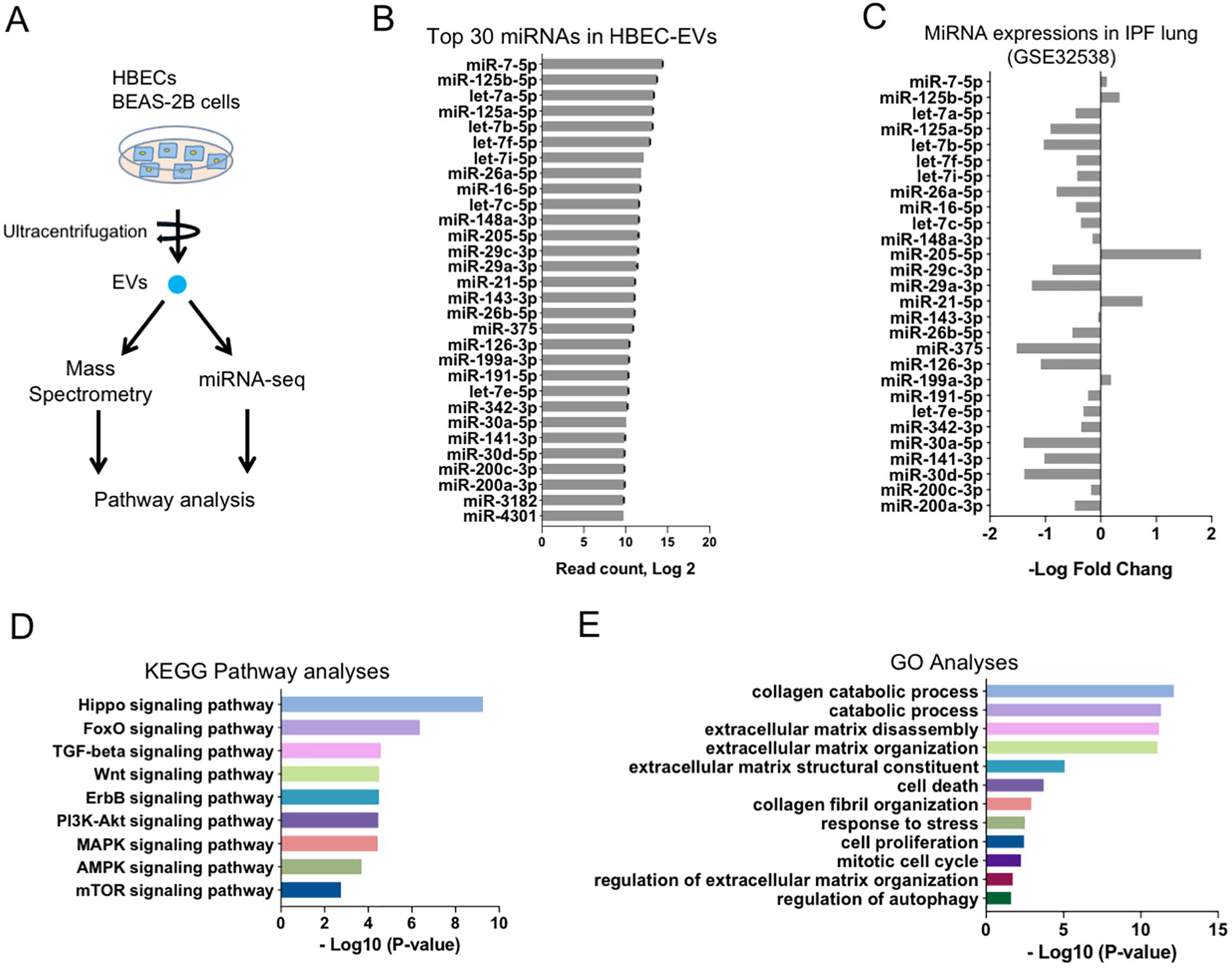
HBEC EVs cargo has a unique miRNA profile. **A** Schematic protocol for the mass spectrometry (MS) and miRNA-seq approach to characterizing EV cargo. EVs were isolated from HBEC and BEAS-2B for MS analysis and miRNA-seq analysis. **B** Summary of the 30 most abundant miRNAs in HBEC EVs. Data represent read counts for each miRNA species in the miRNA-seq analysis. n= 2. **C** Relative expression of the 30 most abundant miRNA species in the miRNA microarray dataset. Bars represent the log 2 fold-change in expression of each miRNA, with positive fold-change values indicating higher levels in IPF lung than non-disease lung. Error bars are omitted for clarity. The Gene Expression Omnibus (GEO) database GSE32538 contains 106 IPF/usual interstitial pneumonitis (UIP) samples and 50 non-diseased samples. **D** Kyoto Encyclopedia of Genes and Genomes (KEGG) pathways from DIANA-mirPath v3.0 analysis, based on the 30 most abundant miRNAs in HBEC EVs. **E** Gene Ontology from DIANA-mirPath v3.0 analysis, based on the 30 most abundant miRNAs in HBEC EVs.

### MiRNA cargo in HBEC EVs inhibits TGF-β-induced expression of WNT5A and WNT10B in LFs

Ingenuity Pathway Analysis (IPA) analysis of RNA-seq data from LFs (Figs. 2C, D) revealed upstream participation of WNT5A, WNT3A, WNT1, and WNT10B in response to the treatment of LFs with HBEC EVs, suggesting that these WNT signaling pathways might represent the major target of HBEC EV miRNA cargo (Fig. 4A). Among these WNT species, only WNT5A and WNT10B are detectable by qRT-PCR in LFs, and expression of these two WNTs are significantly suppressed by HBEC EVs without TGF-β treatment (Fig. 4B). The involvement of WNT5A and WNT10B was further supported by the ability of these recombinant WNT ligands to induce myofibroblast differentiation that was then suppressed by HBEC EVs (Fig. 4C). Previous papers have reported the participation of WNT signaling pathways in IPF pathogenesis (4, 5), and mRNA microarray data sets of GSE53845 and GES10667 from the GEO database reveal significantly increased expression levels of both WNT5A and WNT10B transcripts in IPF lungs (Figs. 4D, E, and Fig. S4A). Considering the possible clinical relevance of anti-fibrotic HBEC EVs for IPF treatment, it is important to show that selected miRNAs in HBEC EVs can directly suppress WNT5A and WNT10B expression. *In silico* analysis using TargetScan Human 7.2 (www.targetscan.org) allowed us to select 16 miRNAs targeting WNT5A and 19 miRNAs targeting WNT10B. Among the 16 miRNAs targeting WNT5A, miR-26a, miR-26b, miR-141a, and miR-200a are included in the 30 most abundant miRNAs present in HBEC EVs (Fig. 4F and Fig. S4B). Transfection of each these four miRNA mimics efficiently suppressed WNT5A expression (Fig. 4G). Among the 19 mRNAs targeting WNT10B, miR-16, miR-29a, miR-29c, and miR-148a are included in top 30 miRNAs present in HBEC EVs (Fig. 4F and Fig. S4C). However, only the miR-16 and miR-148a mimics transfection significantly suppressed WNT10B expression (Fig. 4H). The anti-fibrotic properties of miR-16, miR-26a, miR-26b, miR-141, miR-148a, and miR-200a are demonstrated by their ability to suppress TGF-β-induced myofibroblast differentiation, as judged by type I collagen and αSMA expression (Fig. 4I). Furthermore, expression levels of candidate miRNAs were significantly increased in LFs after treatment with HBEC EVs, indicating the efficient transfer of these miRNAs species into recipient LFs (Fig. 4J).

**Figure 4.**
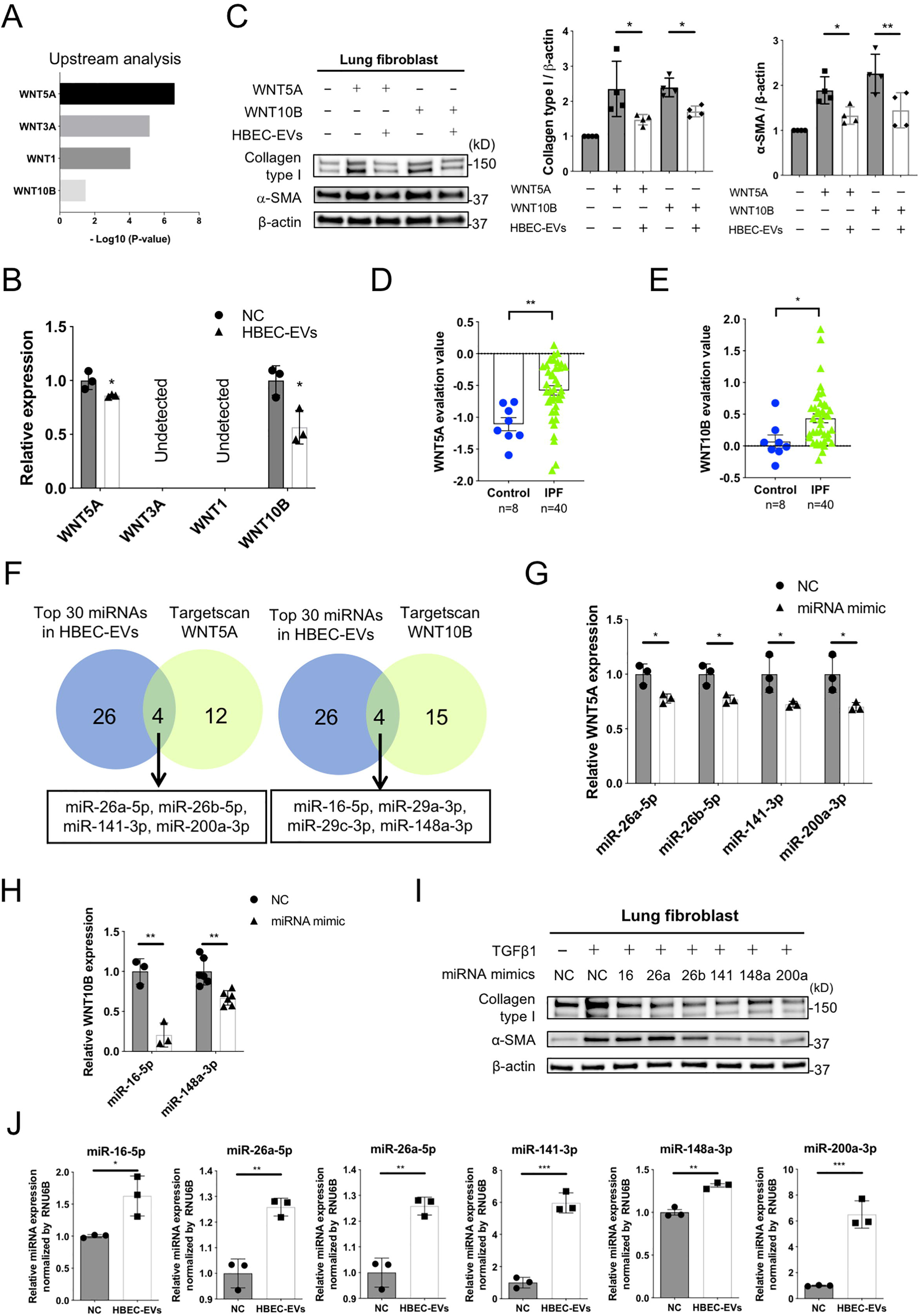
miRNA cargo in HBEC EVs attenuates TGF-β-induced myofibroblast differentiation via suppression of *WNT5A* and *WNT10B expression*. **A** Ingenuity Pathway Analysis (IPA) of upstream regulators of WNT ligand genes in LFs treated for 48 h with HBEC EVs. **B** qRT-PCR analysis of *WNT1, WNT3A, WNT5A* and *WNT10B* mRNAs in LFs treated for 24h with HBEC EVs (10 μg/ml, n=3). \**P*<0.05. **C** Representative immunoblot and quantitative analysis showing the amount of type I collagen, α-SMA, and β-actin in LFs treated for 24 h with PBS or HBEC EVs (10 μg/ml) in the presence or absence of WNT5A (200 pg/ml, lane 2,3) and WNT10B (200 pg/ml, lane 4,5). Protein samples were collected after 72 h of stimulation. *p<0.05, **p<0.005. **D, E** WNT5A and WNT10B evaluation value in the mRNA microarray dataset GSE53845 from the Gene Expression Omnibus (GEO) database. *p<0.05, **p<0.005. **F** Venn diagram of the candidate miRNAs targeting *WNT5A* and *WNT10B* from TargetScan database. **G, H** qRT-PCR analysis of *WNT5A* and *WNT10B* mRNAs in LFs transfected with candidate target miRNA mimics or negative control. \**P*<0.05, **p<0.005. **I** Representative immunoblot analysis showing the amount of type I collagen, α-SMA, and β-actin in LFs treated with validated miRNA mimics or negative control in the presence or absence of TGF-β1 (2 ng/ml). **J** qRT-PCR analysis of cellular miRNA expression in LFs incubated for 24 h with HBEC EVs (10 μg/ml, n=3) or PBS. *p<0.05, **p<0.005, ***p<0.001.

### miRNA cargo in HBEC EVs inhibits TGF-β-induced WNT3A, WNT5A, and WNT10B expressions in HBECs

To implicate miRNA-mediated WNT regulation as a player in the anti-senescence properties of HBEC EVs, we examined the effect of HBEC EVs on TGF-β-induced β-catenin activation in HBECs. In line with the results using LFs, HBEC EVs clearly suppressed TGF-β-induced β-catenin activation in HBECs, with concomitantly reduced expression of p21 as a read-out for senescence (Fig. 5A). Among candidate WNT ligands, WNT5A, WNT3A, and WNT10B were detected in HBECs, all of which were significantly reduced by HBEC EVs treatment (Fig. 5B). Participation of WNT3A, WNT5A, and WNT10B in HBEC EV-mediated suppression of cellular senescence was further evaluated using recombinant WNT ligands. WNT3A, WNT5A, and WNT10B clearly induced p21 expression and cellular senescence in HBECs (Figs. 5C, D, Fig. S5A). HBEC EVs significantly inhibited the increase in p21 expression levels induced by WNT3A, WNT5A and WNT10B and the number of SA-β-positive cells induced by WNT3A and WNT5A. These results suggest that HBEC EV-mediated suppression of cellular senescence is primarily mediated via inhibition of WNT3A and WNT5A (Figs. 5C, D, Fig. S5A). Based on *in silico* analysis using TargetScan, 17 miRNAs targeting WNT3A were selected, of which only miR-16 is included among highly expressed miRNAs in HBEC EVs (Fig. 5E, Fig. S5B). Transfection with miR-16 mimics efficiently suppressed TGF-β-induced WNT3A mRNA expression in HBECs (Fig. S5F). Consistent with data from LFs, miR-26a, miR-26b, miR-141, and miR-200a mimics significantly suppressed WNT5A mRNA levels in HBECs (Fig. S5C). miR-16, miR-26a, miR-26b, miR-141, and miR-200a mimics suppressed TGF-β-induced p21 expression and the percentage of cells staining positive for SA-β-gal (Figs. 5G, H, Fig. S5D). Taken together, these findings suggest the likelihood that these specific cargo miRNA present in HBEC EVs are responsible for suppressing not only myofibroblast differentiation, but also HBEC senescence via regulation of WNT signaling pathways.

**Figure 5.**
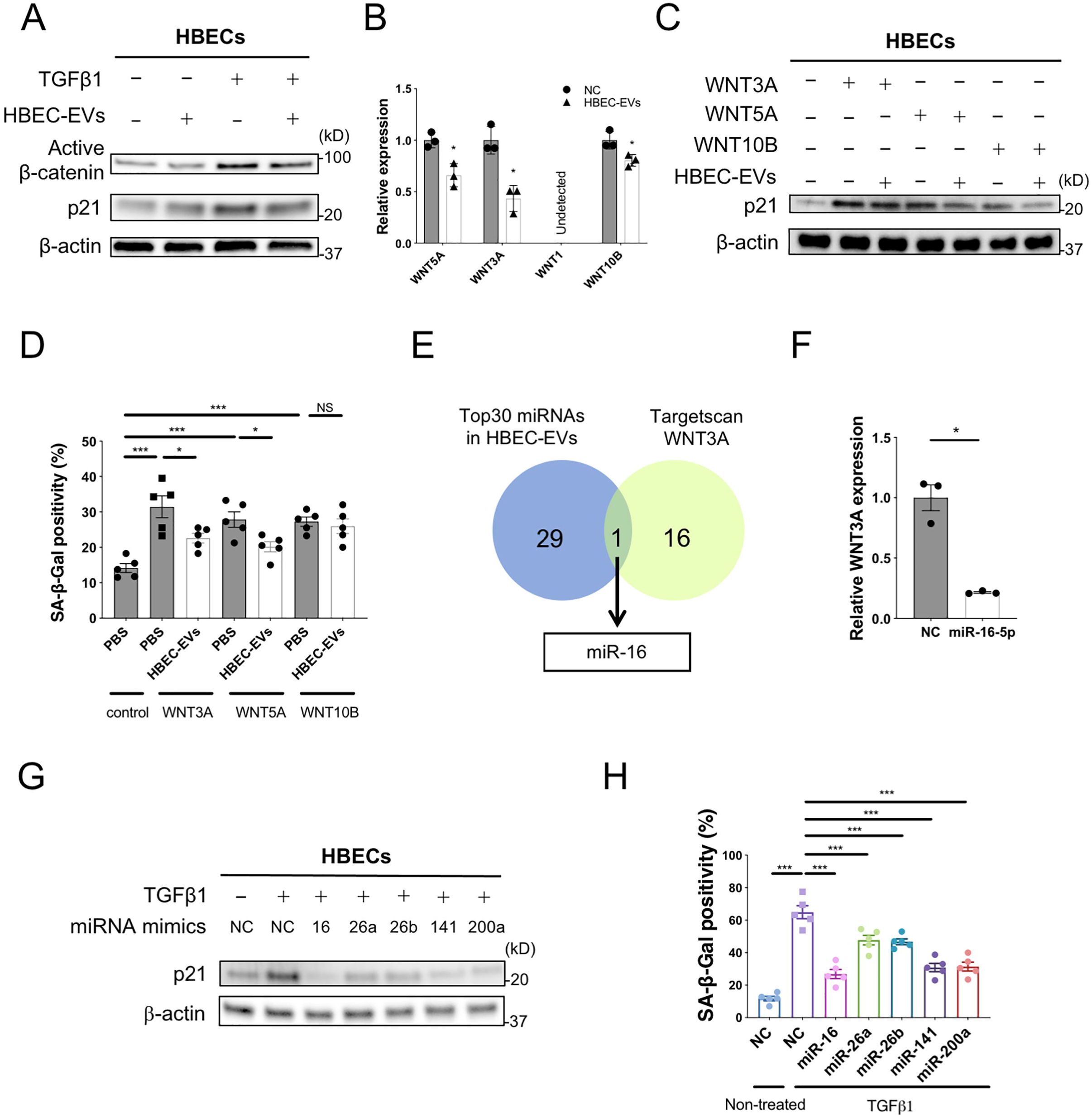
miRNA cargo in HBEC EVs attenuates TGF-β-induced epithelial cell senescence via suppression of *WNT3A* and *WNT5A expression*. **A** Representative immunoblot showing the amount of active β-catenin and β-actin in HBECs treated for 48 h with PBS or HBEC EVs (10 μg/ml) in the presence or absence of TGF-β1 (2 ng/ml). **B** qRT-PCR analysis of *WNT1, WNT3A, WNT5A* and *WNT10B* mRNAs in HBECs treated for 24 h with HBEC EVs (10 μg/ml, n=3). \**P*<0.05. **C** Representative immunoblot analysis showing the amount of p21 and β-actin in HBECs treated for 48 h with PBS or HBEC EVs (10 μg/ml) in the presence or absence of WNT3A (200 pg/ml, lane 2,3), WNT5A (400 pg/ml, lane 4,5), or WNT10B (200 pg/ml, lane 6,7). Protein samples were collected 96 h after stimulation. **D** The percentage of SA-β-gal positive HBECs after treatment for 48 h with PBS or HBEC EVs (10 μg/ml) in the presence or absence of WNT3A (200 pg/ml), WNT5A (400 pg/ml), or WNT10B (200 pg/ml). Staining was performed 96 h after stimulation. * *P*<0.05, \*\*\**P*<0.001. NS; not significant. **E** Venn diagram of the candidate miRNAs targeting *WNT3A* from TargetScan database. **F** RT-PCR analysis of *WNT3A* mRNAs in HBECs transfected with miR-16 mimics or negative control. \**P*<0.05. **G** Representative immunoblot analysis showing the amount of p21 and β-actin in HBECs treated with validated miRNA mimics or negative control in the presence or absence of TGF-β1 (2 ng/ml). **H** The percentage of SA-β-gal positive HBECs after treatment with validated miRNA mimics or negative control in the presence or absence of TGF-β1 (2 ng/ml). \*\*\**P*<0.001.

### Intratracheal administration of HBEC EVs attenuates bleomycin-induced lung fibrosis

We used a mouse model of BLM-induced lung fibrosis to further probe the physiological anti-fibrotic properties of HBEC EVs during the development of lung fibrosis. We followed the lead of a previous paper in which 2 × 10^9^ HBEC EVs or BM-MSC EVs were used as a dose for intratracheal administration (28). Intratracheal injection was performed at days 7 and 14 after BLM treatment to evaluate the anti-fibrotic effects of EVs on the progression of fibrosis (Fig. 6A). In general, day 7 is considered to be the beginning of the fibrotic phase in this model, along with concomitant resolution of the acute inflammatory reaction (29). Efficient uptake and delivery of EVs to the distal lungs were confirmed by using HBEC EVs labelled with PKH67 green fluorescence (Fig. S6A). Compared to control mice, significant loss of body weight was detected in BLM-treated mice, a deficit that was rescued by HBEC EV treatment (Fig. 6B). Intratracheal administration of both HBEC EVs and BM-MSC EVs significantly attenuated the development of BLM-induced lung fibrosis, as evaluated by Achcroft score (Fig. 6C) (30). Attenuation of the development of BLM-induced lung fibrosis was further confirmed by both Masson trichrome staining and hydroxyproline assay (Figs. 6D, E). In line with *in vitro* experiments, HBEC EVs exerted more potent anti-fibrotic effects on BLM-induced lung fibrosis than seen with MSC EVs (Figs. 6C, D, E). The importance of HBEC EV-mediated suppression of WNT signaling in the BLM model was confirmed by the use of immunohistochemical staining to demonstrate reduced levels of β-catenin expression in EV-treated lungs (Fig. 6F). BLM-induced expression of the senescence markers p16 and p21 was clearly suppressed by HBEC EVs treatment (Fig. 6F). Importantly, no adverse effects of intratracheal EV administration were detected in post-mortem serological examinations, indicating the safety of the procedure in the context of clinical application (Fig. S6B).

**Figure 6.**
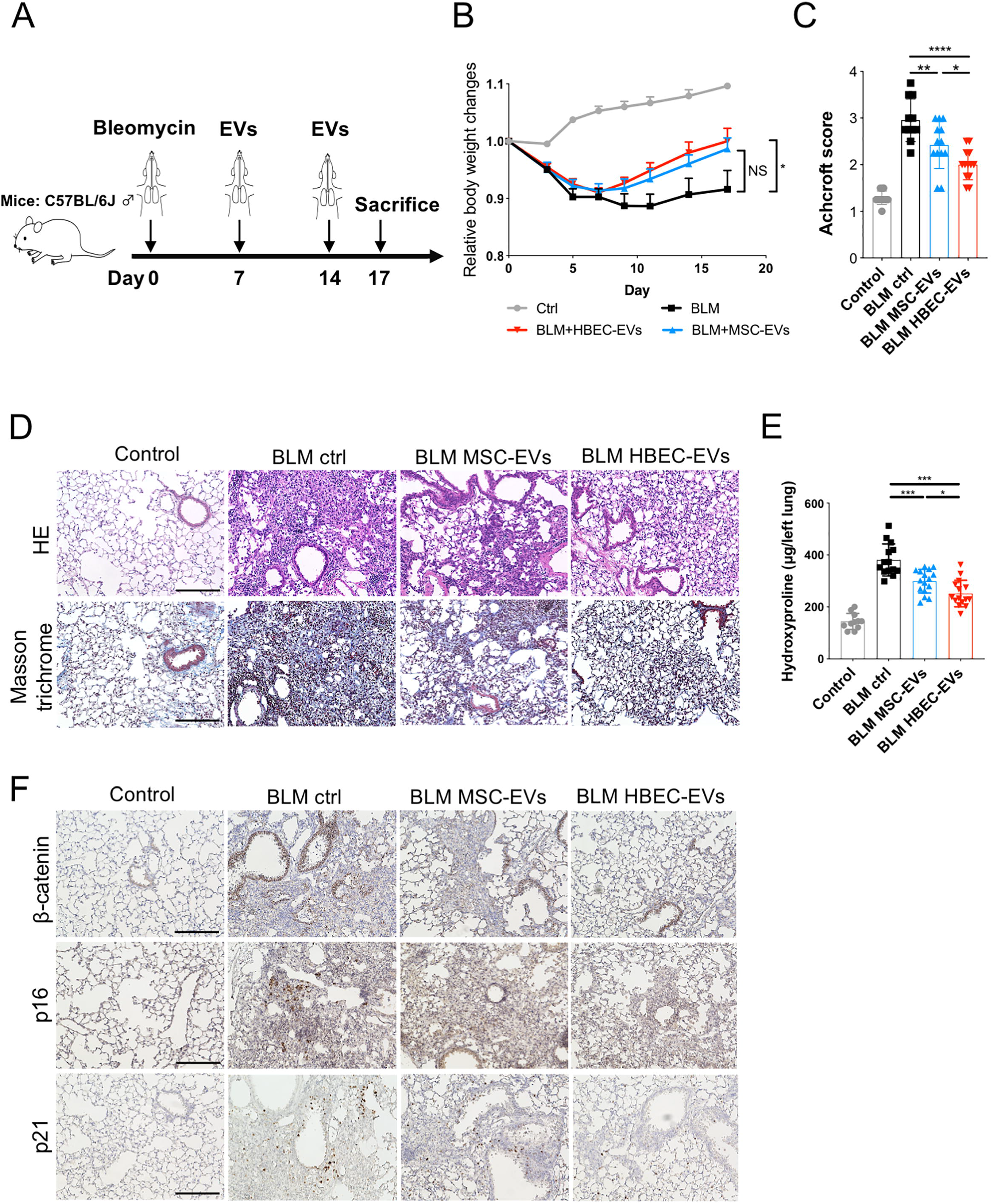
Intratracheal administration of HBEC EVs attenuates bleomycin-induced lung fibrosis in mice. **A** Schematic protocol for EV treatment in the mouse of bleomycin-induced pulmonary fibrosis. **B** Body weight (BW) changes after bleomycin treatment. BW at d 0 before initiating treatment was given a relative value of 1.0. Graphed values represent the mean ± SEM. \**P*<0.05. Control n = 11, BLM ctrl n = 8, BLM MSC EVs n = 10, BLM HBEC EVs n = 11. **C** Quantification of fibrosis by Ashcroft score. Each dot represents data from one animal. \**P*<0.05, \*\**P*<0.005, \*\*\*\**P*<0.0005. Control n = 12, BLM ctrl n = 11, BLM MSC EVs n = 13, BLM HBEC EVs n = 13. **D** Masson’s trichrome and Hematoxylin and Eosin staining of representative lung sections (n=11) from each indicated group of treated mice. Scale bars, 200 μm. **E** Hydroxyproline content from each indicated group of treated mice. Each dot represents data from one animal. \**P*<0.05, \*\*\**P*<0.001. Control n = 11, BLM ctrl n = 14, BLM MSC EVs n = 16, BLM HBEC EVs n = 15. **F** Immunohistochemical staining of β-catenin, p21 and p16 in representative lung sections (n=11) from each indicated group of treated mice. Scale bars, 200 μm.

## Discussion

MSC cell therapy is proposed to be a potentially effective modality of treatment for a number of refractory disorders, including IPF (31, 32). ATII dysfunction with concomitantly increased cell loss conferred due to regulated cell death (RCD) has been widely implicated in IPF pathogenesis (33, 34). ATII is recognized to be a progenitor cell with capacity to proliferate and differentiate into alveolar type I cells (ATI) during the process of alveolar wound healing. Therefore, homing to sites of lung injury and differentiating to ATII are proposed to represent underlying mechanisms for the efficacy of MSC cell therapy. Although the safety of intravenous MSC injection was demonstrated in IPF patients in a phase I clinical trial, it has not been completely excluded that MSCs might be the potential pathogenic source of abnormal fibroblasts or that MSCs might possess malignant potential without the ability to effectively restore the ATII population (9, 35). In addition, clinical application of cell injection therapies has been fraught with technical challenges in terms of not only efficacy but also scalability, reproducibility, and safety. As an alternative, it has been reported that EVs may carry cargo that endows them with properties similar to those of parental cells, and indeed, that MSC EVs have therapeutic effects on regeneration and immunomodulation (12, 15, 17). Intravenous administration of MSC EVs has produced attenuation of BLM-induced lung fibrosis via modulation of modulating monocyte phenotypes (18). Thus, EVs may serve as an alternative to cell therapies. In the context of hypoxia-induced lung injury model, intratracheal administration of MSCs has been shown to exert a more protective effect than systemic intravenous administration, suggesting the possible advantage of inhalation treatment (36–38). Importantly, with respect to developing inhalation therapy as a specific treatment modality for respiratory diseases, EVs may have several advantages for clinical application. First, the smaller size of EVs enhances their delivery and deposition in the small airway and alveolar regions responsible for IPF development. Second, EVs are stable in tissues and body fluids due to their lipid bilayer architecture. Third, compared to cell therapy, EVs exhibit low levels of immunogenicity and toxicity. Based on these advantages, intratracheal administration of EVs may represent an ideal treatment modality for refractory lung disorders. In fact, a recent paper shows that inhalation of lung spheroid cell-derived exosomes efficiently attenuates BLM-induced lung fibrosis by decreasing myofibroblast proliferation (19).

In the present study, we demonstrate that HBEC EVs efficiently suppress TGF-β-induced myofibroblast differentiation and lung epithelial cell senescence, and that the effects of HBEC EVs are more potent than those observed with MSC EVs. Based on RNA-seq analysis, we show that both canonical and non-canonical WNT signaling pathways are involved in the mechanisms by which HBEC EVs mediate suppression of TGF-β-induced myofibroblast differentiation and cellular senescence. KEGG pathway analysis and GO analysis reveal that miRNA cargo in HBEC EVs is responsible for attenuating signaling by TGF-β and WNT. Among enriched miRNA species present in HBEC EVs, miR-26a, miR-26b, miR-141, and miR-200a target non-canonical WNT5A in both LFs and HBECs. MiR-16 and miR-148a target canonical WNT10B in LFs, and miR-16 targets canonical WNT3A in HBECs. Compared to MSC EVs, intratracheal administration of HBEC EVs, in the context of a mouse model of BLM-induced lung fibrosis, more potently attenuates cellular senescence and lung fibrosis, effects that are accompanied by reduced β-catenin expression. HBEC EVs significantly suppress TGF-β-induced myofibroblast differentiation and cellular senescence in IPF LFs. These findings indicate that HBEC EVs may possess therapeutic anti-fibrotic properties that are effective in preventing development of lung fibrosis in IPF.

Activation and cooperation between TGF-β and WNT signaling pathways are important for the normal developmental processes of airway abnormal wound healing process that occurs during fibrotic remodeling in IPF (4, 5). Aberrant recapitulation of this pattern of developmental gene expression has been widely proposed to be implicated in the TGF-β plays pivotal roles in lung fibrosis via mediation of myofibroblast differentiation, epithelial-mesenchymal transition (EMT), cellular senescence, and epithelial cell apoptosis (23, 25, 39). Both β-catenin dependent (canonical) and β-catenin independent (non-canonical) WNT signaling pathways are responsible for epithelial cell proliferation, cellular senescence, EMT, and myofibroblast differentiation (5, 40–44). Since TGF-β and WNT have overlapping roles in regulating cell phenotype and cell fate, it is likely that functional crosstalk between TGF-β and WNT signaling pathways play an essential role in IPF pathogenesis (24, 41). Hence, TGF-β-WNT crosstalk has been proposed to be a promising target for treatment of IPF (5, 45). We have confirmed the involvement of WNT signaling pathways in the mechanisms underlying HBEC EV-mediated suppression of TGF-β-induced myofibroblast differentiation and cellular senescence. This involvement occurs via the reduction of WNT ligand expression by targeting with miR-16, miR-26a, miR-26b, miR-141, miR-148a, and miR-200a. This mechanism is supported by results of a previous paper showing significant reduction of miR-26a concomitant with activation of TGF-β signaling in IPF lungs (46). In addition, attenuation of experimental lung fibrosis was achived by overexpressing miR-26a (46). It has also been reported that the miR-200 family can inhibit TGF-β1-induced EMT of alveolar epithelial cells, but that this miRNA family is downregulated in IPF (47, 48). Reduced cellular senescence in ATII was also archived via miR-200 and miR-141 treatment (48). Collectively, it is not surprising that selected miRNAs such as miR-26a, miR-200a, and miR-141 are responsible for inhibiting TGF-β-WNT crosstalk and are involved in the antifibrotic properties of HBEC EVs. Among other miRNAs abundantly represented in HBEC EVs, miR-29 is down-regulated in BLM-induced fibrotic lungs, and a negative correlation exists between miR-29 expression and genes for fibrotic proteins such as *COL1A1, COL1A2, FBN1* expression (46, 49). In another example, reduction of *Let-7* family is detected in IPF lungs and *let-7d* is down regulated by TGF-β via SMAD3 binding to the *let-7d* promoter (50). The pro-fibrotic effect of *let-7d* down-regulation was seen in both *in vitro* and *in vivo* experiments (50). Intriguingly, a recent paper demonstrates that reduced *let-7* is associated with pulmonary fibrosis through expression of the estrogen receptor (ER) (51). Therefore, it is plausible that high expression of these miRNAs can attributed to the anti-fibrotic effect of HBEC EVs through different mechanisms of regulating TGF-β-WNT crosstalk. Upregulation of pro-fibrotic miR-21 in IPF lungs is known to promote accumulation of myofibroblasts via activating TGF-β signaling pathway (52). In our study, miR-21 is one of the abundant miRNAs in HBEC EVs, suggesting the possibility that pro-fibrotic miRNAs such as miR-21 may interfere with the anti-fibrotic capacity of HBEC EVs (Fig. 3B). Nevertheless, KEGG pathway analysis of 30 highly expressed EV miRNAs including miR-21 clearly demonstrate the participation of EVs in suppressing TGF-β and WNT signaling. We therefore consider that overall, the composite miRNA cargo in HBEC EVs serves to attenuate lung fibrosis via regulation of TGF-β-WNT crosstalk for clinical translation.

It is interesting to speculate about the possible role of EVs in normal lung physiology, especially in the context of airway-alveolar interaction. Based on the physical properties of EVs, we suppose that HBEC EVs may be transferred to both epithelial cells and fibroblasts in alveoli as a means of maintaining homeostasis of the microenvironment in healthy lungs (Fig. S6C). On the other hand, as demonstrated by CS-exposed HBEC EVs (21), it is plausible that EVs derived from metaplastic epithelial cells (ME) during bronchiolization may have a pathological profibrotic role during the aberrant wound healing process in IPF. This pathology might be controlled by administration of sufficient numbers of physiological HBEC EVs as a means of regulating TGF-β and WNT signaling pathways (Fig. S6C).

This issue points to one of several limitations in our present study. First, it is plausible that other non-miRNAs components carried by HBEC EVs may participate in anti-fibrotic mechanisms. Further assessment of this possibility should be made. Second, due to the pleiotropic role of TGF-β in targeting a wide array of downstream regulators, the mechanism underlying HBEC EV-mediated inhibition of TGF-β action is not restricted to WNT crosstalk. Other mechanisms involving TGF-β should also be investigated. Third, TGF-β is not the only driver of lung fibrosis in IPF. Other anti-fibrotic mechanisms of HBEC EV action, besides regulation of TGF-β-WNT crosstalk, should therefore be examined. Fourth, although accelerated cellular senescence in metaplastic epithelial cells is a pathogenic hallmark of IPF lungs (25), technical restriction of the cell culture system did not allow us to determine the anti-senescence property of HBEC EVs with respect to alveolar epithelial cells. Fifth, because of limitations in IPF sample collection, the clinical implication of HBEC EVs was evaluated only in a limited number of IPF LFs, and we could not examine anti-senescence phenomena using IPF lung epithelial cells. Accordingly, it is critically important prior to clinical application that the detailed pharmacotherapeutic properties of HBEC EVs should be further examined using IPF samples.

In conclusion, we show that the anti-fibrotic properties of normal HBEC EVs are more potent than those of MSC EVs and that this effect is at least partly mediated by miRNA cargo via negative regulation of TGF-β-WNT crosstalk. We observed efficient attenuation not only of the profibrotic cell phenotype and experimental lung fibrosis in vivo, but also myofibroblast differentiation and cellular senescence in IPF LFs. These findings indicate that intratracheal administration of HBEC EVs can be a promising cell-free antifibrotic modality of treatment for IPF via targeting of TGF-β-WNT crosstalk. Treatment safety and efficacy necessary for clinical translation will be examined in future studies.

## Materials and Methods

### Cell culture and clinical samples

All human lung tissues were obtained from pneumonectomy and lobectomy specimens at The Jikei University School of Medicine. HBECs were purchased from Lonza or were isolated from normal airway tissue (1st through 4th order bronchi) using protease treatment as previously described (53). Freshly isolated HBECs were plated onto rat tail collagen type I-coated (10 μg/ml) dishes in Bronchial Epithelial Growth Medium (BEGM, Lonza) containing 1% antibiotic– antimycotic solution (AA, Invitrogen) and grown at 37 °C in 5% CO_2_. HBECs were used up to passage 3. The bronchial epithelial cell line BEAS-2B was cultured at 37 °C in 5% CO_2_ in Roswell Park Memorial Institute (RPMI) 1640 medium containing 10% fetal bovine serum (FBS, Gibco Life Technologies, 26140-079) and supplemented with a 1% AA. LFs were isolated and characterized using an explantation technique. Briefly, LFs growing out from lung fragments were cultured in Dulbecco’s modified Eagle’s medium containing 10% FBS and 1% AA. LFs were isolated and characterized as previously described (54) and summarized here. During the process of normal LF isolation, histological evaluation of lung tissue was performed to exclude the inclusion of cancer tissues and ensure fibroblasts were derived only from normal tissues. LFs derived by outgrown from lung fragments were cultured in fibroblast growth medium (DMEM containing 10% FCS and 1% AA). LFs were serially passaged when the cells reached approximately 80% to 100% confluence and were still actively proliferating. LFs were used for experiments up to passage 6. HSAECs were isolated from distal portions of lung tissue. Lung tissue fragments were mechanically minced and dissociated with enzymes according to a Lung Dissociation Kit protocol (Miltenyi Biotec, 130-095-927). A Miltenyi Gentle MACS dissociator was used for mincing, followed by incubation with enzymes for 42 min at 37 °C. Cells were then washed and passed through 70-μM and 40-μM filters. The dissociated cell suspension was cultured on rat tail collagen type I-coated dishes at 37 °C in 5% CO_2_ using Small Airway Epithelial Cell Growth Medium (SAGM; basal medium plus growth supplements, Lonza, CC-3118) containing 1% AA. Human BM-MSCs were purchased from the RIKEN cell bank. BM-MSCs were routinely cultured in reduced serum (2%) medium (MesenPRO RS, Invitrogen) containing 1% Glutamax (Invitrogen) and 1% AA. BM-MSCs were used for experiments within ten passages. Informed consent was obtained from all surgical participants as part of an ongoing research protocol approved by the Ethical Committee of The Jikei University School of Medicine (#20-153 (5443)). The diagnosis of IPF was based on an international consensus statement from the American Thoracic Society and the European Respiratory Society (55). High-resolution chest CT images were used to fulfill the criteria for recognizing UIP pattern. Secondary ILD samples such as connective tissue disease-associated, hypersensitivity pneumonitis, radiation treatment, drug-induced, occupation-associated and sarcoidosis were excluded. Control participants had no history of pulmonary fibrosis, COPD or any other inflammatory diseases.

### Antibodies and reagents

Antibodies used were rabbit anti-p21 (Cell Signaling Technology, 2947), rabbit anti-p16 (Santa Cruz Biotechnology, sc-468), mouse anti-β-actin (Millipore, Billerica, MAB1501), mouse anti–α smooth muscle actin (Sigma-Aldrich, A2547), goat anti-type I collagen (SouthernBiotech, 1310-01), rabbit anti-β-catenin (Cell Signaling Technology, 8480), rabbit anti-non-phospho (active) β-catenin (Cell Signaling Technology, 8814), rabbit anti-histone H3 (Cell Signaling Technology, 4499), mouse anti-CD81 (Santa Cruz Biotechnology, 555675), mouse anti-CD9 (Santa Cruz Biotechnology, sc-59140), mouse anti-CD63 (BD Pharmingen, 556019), rabbit anti-Hsc70 antibody (Proteintech, 10654-1-AP) and mouse anti– Tom20 (Santa Cruz Biotechnology, sc-17764). Hoechst 33242 (Sigma-Aldrich, H342) and collagen type I solution from rat tail (Sigma-Aldrich, C3867) were purchased reagents.

### EV purification and analysis

Following growth to 50-80% confluence on 10 cm-cell culture dishes, cells for EV preparation (HBECs, BEAS-2B, HSAECs and BM-MSCs) were washed with PBS. Culture medium was then replaced with fresh BEGM containing 1% AA (for HBECs), advanced RPMI 1640 containing 1% AA and 2 mM L-glutamine (for BEAS-2B), SAGM containing 1% AA (for HSAECs), or MSC serum-free medium (SFM, Gibco) containing 1% AA (for BM-MSCs). After incubation for 48 h, conditioned medium (CM) was collected and centrifuged at 2,000 g for 10 min at 4 °C. The supernatant was then passed through a 0.22 μm filter (Millipore) to thoroughly remove cellular debris. For EV preparation, the CM was ultracentrifuged at 4 °C for either 70 min at 35,000 r.p.m. using a SW41Ti rotor or for 45 min at 44,200 r.p.m. using a MLS50 rotor. Resulting pellets were washed by ultracentrifugation in PBS and then resuspended in PBS. Protein concentrations of the putative EV fractions were measured using a Quant-iT Protein Assay and a Qubit 2.0 Fluorometer (Invitrogen). To determine the size distribution of the EVs, nanoparticle tracking analysis was carried out using the Nanosight system (NanoSight) with samples diluted 500-fold with PBS for analysis. This system focuses a laser beam through a suspension of the particles of interest. Particles are visualized by light scattering using a conventional optical microscope perpendicularly aligned to the beam axis, allowing collection of light scattered from every particle in the field of view. A 60 s video recording of all events is then used for further analysis by nanoparticle tracking software. The Brownian motion of each particle is used to calculate its size via the Stokes–Einstein equation.

### RNA extraction and qRT-PCR

Total RNA was extracted from cultured cells or EVs using QIAzol (Qiagen) and the miRNeasy Mini Kit (Qiagen) in accordance with the manufacturer’s protocol. For mRNA expression by qRT–PCR analysis, complementary DNA (cDNA) was generated from total RNA using a High capacity cDNA Reverse Transcription Kit (Thermo Fisher Scientific). For miRNA expression by qRT–PCR analysis, cDNA was generated from total RNA using a TaqMan microRNA Reverse Transcription Kit (Thermo Fisher Scientific). Real-time PCR was subsequently performed with cDNA using a universal PCR Master Mix (Thermo Fisher Scientific). The data were collected and analyzed using a StepOne Real-Time PCR System and StepOne Software v2.3 (Thermo Fisher Scientific). All mRNA quantification data from cultured cells were normalized to the expression of β-actin. All miRNA quantification data were normalized to the RNUB6. All pcr primers were purchased from Thermo Fisher Scientific and are listed in Supplementary Table 4 and Table 5.

### Cell transfection

Transfection of miRNA species at a concentration of 30 nM was conducted with Lipofectamine RNAiMAX reagent (Thermo Scientific) according to the manufacturer’s protocol. LFs and HBECs to be transfected were seeded at 50-70 % confluence the day before transfection. MiR-16-5p (MC 10339, MH10339), miR-26a-5p (MC10249, MH10249), miR-26b-5p (MC12899, MMH12899), miR-141-3p (MC10860), miR-148-3p (MC 10263), miR-200a-3p (MC10991) and negative control miRNA (Ambion) were used for these transfections.

### Subcellular fractionation and Immunoblot analysis

Nuclear and cytoplasmic fractions were prepared using the Nuclear and Cytoplasmic Extraction reagents (Thermo-Fisher Scientific, 78833) according to the manufacturer’s instructions. HBECs were lysed in Mammalian Protein Extract Reagent (M-PER; Thermo Scientific, Rockford, 78501) with sample buffer solution (Wako, 198-13282 or 191-13272). For each immunoblotting experiment, equal amounts of total protein were loaded onto 4–20% SDS-PAGE gels (Mini-PROTEAN TGX Gel, Bio-Rad). After SDS-PAGE, proteins were transferred to a polyvinylidene difluoride membrane (Millipore). After blocking in Blocking One (Nacalai Tesque, Kyoto, Japan), the membranes were incubated for 1 h at room temperature with specific primary Ab. Anti-rabbit IgG (Cell Signaling Technology, 5127), and anti-mouse IgG (GE Healthcare, NA931) were used as secondary antibodies at 1:1,000 dilution. Anti-goat IgG (Jackson ImmunoResearch, 705-035-147) was used at 1:10,000 dilution. Membrane were then exposed to ImmunoStar LD (Wako, Osaka, Japan) with a LAS-4000 UV mini system (Fujifilm, Tokyo, Japan).

### Immunofluorescence staining

After washing three times with PBS, cells were fixed in 4% paraformaldehyde (Wako), and incubated for 1 hour with primary antibodies 0.1% BSA. The cells were then incubated with Alexa Fluor fluorescent secondary antibodies containing 0.1% BSA. DAPI staining was performed immediately before imaging. All staining was observed using a confocal microscope (LSM 880, ZEISS).

### SA-β-Gal Staining

Senescence associated β-galactosidase (SA-β-Gal) staining was performed using HBECs and LFs grown on 24-well culture plates according to the manufacturer’s instructions (Sigma, CS0030).

### Next generation sequencing and bioinformatics

RNA quantity and quality were determined using a NanoDrop ND-1000 spectrophotometer (Thermo Fisher Scientific Inc.) and an Agilent Bioanalyzer (Agilent Technologies). To detect miRNAs present in EVs for next-generation sequencing, the sequencing library was prepared using the QIAseq miRNA Library Kit (Qiagen, 331502), which tags each individual miRNA molecule with a unique molecular index (UMI). This is used in data analysis to correct for library amplification bias, followed by amplification of each miRNA via cDNA synthesis/PCR. Five μl quantities were used for conversion into libraries. Each sample was read an average of 10 million times. Amplified miRNAs were sequenced using the Next-Generation System (Illumina NextSeq 500). Raw data was converted to FASTQ files and checked using the FastQC tool. Reads with identical UMIs were collapsed into a single read and aligned to miRBase Release 21 to identify known miRNAs. StrandNGS was applied to normalize the data using the trimmed mean of M (TMM) method and to determine significant differentially expressed miRNAs between groups. To detect the mRNAs for next-generation sequencing (Illumina HiSeq 2500), the mRNA sequencing library was prepared using theTruSeq Stranded mRNA Sample Prep Kit. Paired-end sequenced reads were checked by FastQC and aligned to the reference genome, human GRCh38, using TopHat2. Cuffdiff was then performed to define FPKM values for genes.

### Mass spectrometric analysis

Protein solution samples were reduced and alkylated as previously described (56). Proteins including EVs were diluted 3-fold with 50 mM Ammonium bicarbonate, pH 8.5, and digested with trypsin at an enzyme-to-substrate ratio of 1:50 (wt/wt, modified sequencing grade, Promega, Madison, Wisconsin. United States) for 16 hours at 30°C. LC-MS/MS analyses were performed by nano LC (UltiMate^®^ 3000) (Dionex, Sunnyvale, CA, USA) coupled with Q Exactive plus orbitrap mass spectrometer (Thermo Scientific, Waltham, MA, USA). Mass spectrometric analysis was performed according to the method described previously (57). The database search was performed with MASCOT Deamon (Matrix Science, London, UK). The generated pkl files were submitted to NCBInr (20130316). Search parameters were as follows: fixed modifications, carbamidomethyl; variable modifications, oxidation (M); missed cleavages, up to 1; monoisotopic peptide tolerance, 1.0 Da; and MS/MS tolerance, 0.5 Da. To eliminate the very-low-scoring random peptide matches automatically, the ions score cutoff was set to 30.

### Animal studies

Animal experiments were performed in compliance with the guidelines of the Institute for Laboratory Animal Research, The Jikei University School of Medicine (Number: 2018-071). C57BL/6J mice (Charles River Laboratories) were used in all experiments. A dose of 2.5 U/kg bleomycin (Nippon Kayaku Co., 4234400D4032) was intratracheally administered in 50 mL saline using the MicroSprayer Aerosolizer and a high-pressure syringe (PennCentury, Philadelphia, PA, USA). HBEC EVs and BM-MSC EVs (2.0 × 10^9^/body) were given intratracheally on day 7 and day 14 after bleomycin administration. Equivalent volumes of 0.9% normal saline were used as controls. On the 17th day, the lung tissue and blood were collected for protein determination, histological examination and blood biochemistry.

### Immunohistochemical staining

Lung tissue samples for immunohistochemistry were collected from C57BL/6J mice. Tissue samples were fixed with formalin and embedded in paraffin. Following dewaxing and rehydration, heat-induced epitope retrieval was performed by boiling the specimens in 1/200 diluted ImmunoSaver (Nissin EM, Tokyo, Japan) at 98 °C for 45 min. Endogenous peroxidase activity was inactivated by treating the specimens with 3% H_2_O_2_ at room temperature for 10 min. Specimens were then treated with 0.1% Triton X-100 for tissue permeabilization. After treatment with a Protein Block Serum-Free blocking reagent (DAKO, Code X0909) at room temperature for 30 min, the specimens were incubated with primary antibodies, anti-rabbit β-Catenin (CST, 8480), anti-mouse p16 (abcam, ab54210), and anti-rabbit p21 (CST, 2947) at room temperature for 1 h or at 4 °C overnight. The sections were then processed using ImPRESS IgG-peroxidase kits (Vector Labs) and a metal-enhanced DAB substrate kit (Life Technologies), according to the supplier’s instructions. Hematoxylin and eosin (HE) and Masson’s trichrome staining were performed on paraffin embedded tissue sections. After counterstaining with haematoxylin, specimens were dehydrated and mounted.

### PKH67-labelled HBEC EVs transfer

Purified HBEC EVs were labelled with a PKH67 green fluorescence labelling kit (Sigma-Aldrich). HBEC EVs (2.0 × 10^9^) were incubated with 2 mM PKH67 for 5 min and then washed five times using a 100 kDa filter (Sartorius, VIVACON 500 100K VN01H41) to remove excess dye. PKH67-labelled EVs were intratracheally administered in 50 μL saline on day 7, and the lungs were collected on the next day.

### Hydroxyproline content of whole lung

For quantitatively measuring collagen in the left lungs of mice, the Hydroxyproline assay was performed according to the manufacturer’s instructions (Chondrex, 6017). Mouse left lung tissues (10mg) were homogenized in 100 μl of distilled water using PowerMasher II (Nippi, 891300) with BioMasherII (Nippi, 320103).

### Blood biochemistry

The blood was collected from C57BL/6J mice in blood anti-coagulant treated centrifuge tubes at day 17 after BLM treatment. All blood analysis was performed according to the International Federation of Clinical Chemistry and Laboratory Medicine (IFCC) using the JCA-BM 8060 instrument from JEOL.

### Statistics and Data analysis

Data presented in bar graphs are shown as the average (±SEM) for technical replicates. A Student t test was used for comparison of two data sets. Analysis of variance (ANOVA) was used for multiple comparisons, followed by Tukey or Dunnett multiple comparison to identify differences. Significance was defined as p< 0.05. Statistical software used was Prism version 7 (GraphPad Software, San Diego, CA). Gene Set Enrichment Analysis (GSEA, www.broadinstitute.org/gsea) was performed on RNA-seq date following EV treatment using gene set collections from the Molecular Signatures database (MSigDB) v6.1. Gene ontology (GO) analyses were performed with Database for Annotation, Visualization and Integrated Discovery (DAVID) v6.8, using all the proteins identified by the EV proteomics experiment as background. DIANA-mirPath v.3.0 software (http://snf-515788.vm.okeanos.grnet.gr/) was performed to predict potential pathways affected by miRNAs present in EVs. The DIANA-microT-CDS target prediction algorithm was employed to identify miRNA targets, which was combined with KEGG and GO databases.

## Data availability

All miRNA and mRNA sequencing data have been deposited in the Gene Expression Omnibus(GEO) (https://www.ncbi.nlm.nih.gov/geo/) database (accession number: GSE156572, GSE158624). The authors declare that the data supporting the findings of this study are available within the paper and its supplementary information file.

## Acknowledgements

We thank Dr. Akihiro Ichikawa, Dr. Yusuke Hosaka and Dr. Akihiko Ito (The Jikei University School of Medicine, Tokyo, Japan) for technical support and clinical sample collection, Misato Yamamoto for technical assistance and Dr. Yusuke Yoshioka (Tokyo Medical University, Tokyo, Japan) for technical advice.

## Author contributions

T.K., Y.F., J.A. and T.Ochiya. conceived and designed the study. T.K. and N.W. performed the experiments. T.K. and Y.F. performed data analysis. T.K., Y.F., J.A., N.W., S.F., H.K., S.M., H.H., Y.Y. and K.K. interpreted the data. T.K,, Y.F. and J.A. wrote the manuscript. T.Otsuka. provided the patient samples. Y.F. and T.Ochiya. supervised this project. All authors reviewed and edited the manuscript.

## Competing Interests statement

The authors have declared that no conflict of interest exists.

## Notes

### Competing Interest Statement

The authors have declared no competing interest.

