## Supplementary figures and images for "Human bronchial epithelial cell-derived extracellular vesicle therapy for pulmonary fibrosis via inhibition of TGF-β-WNT crosstalk"

### Fig. S1

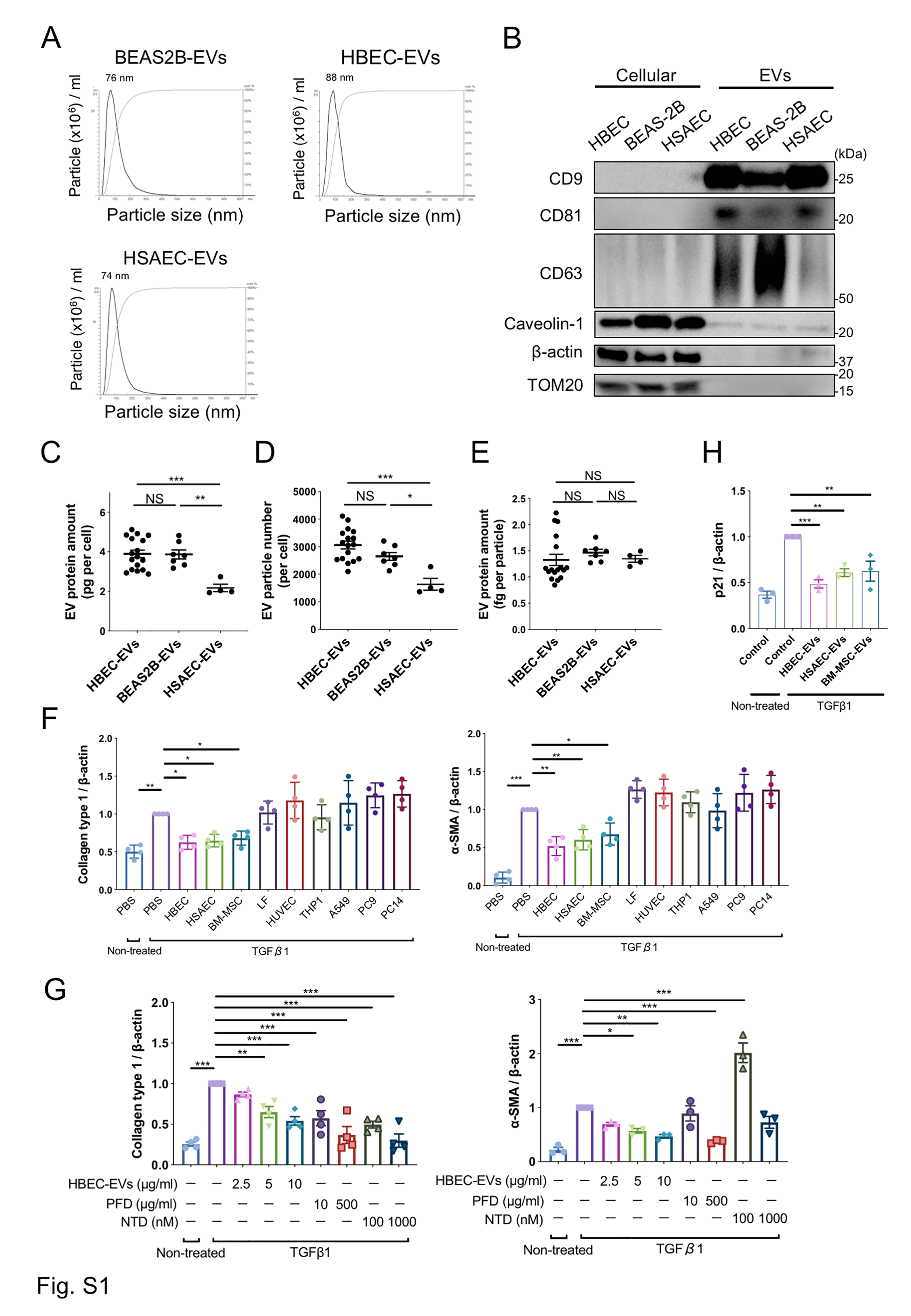

### Fig. S2

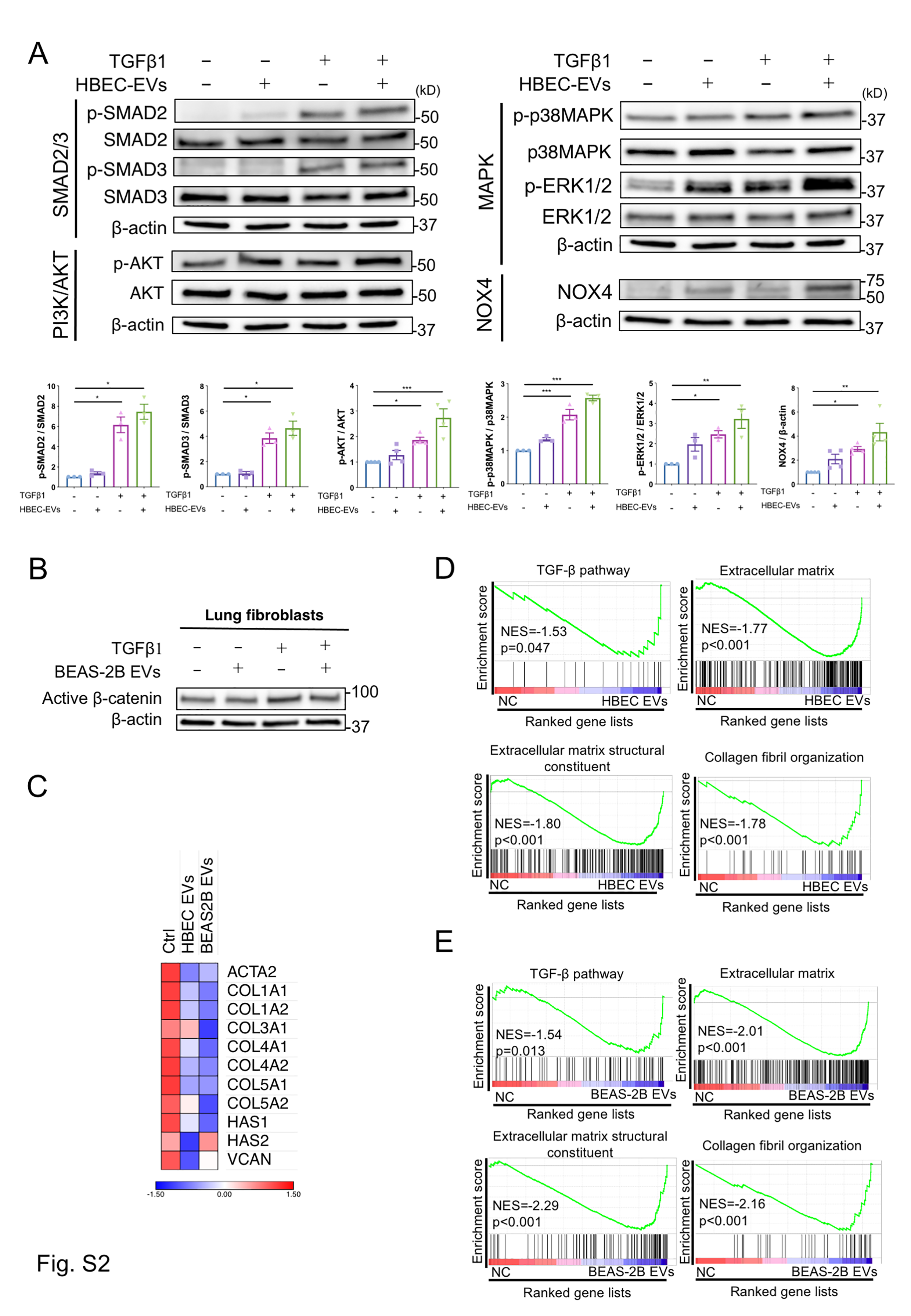

### Fig. S3

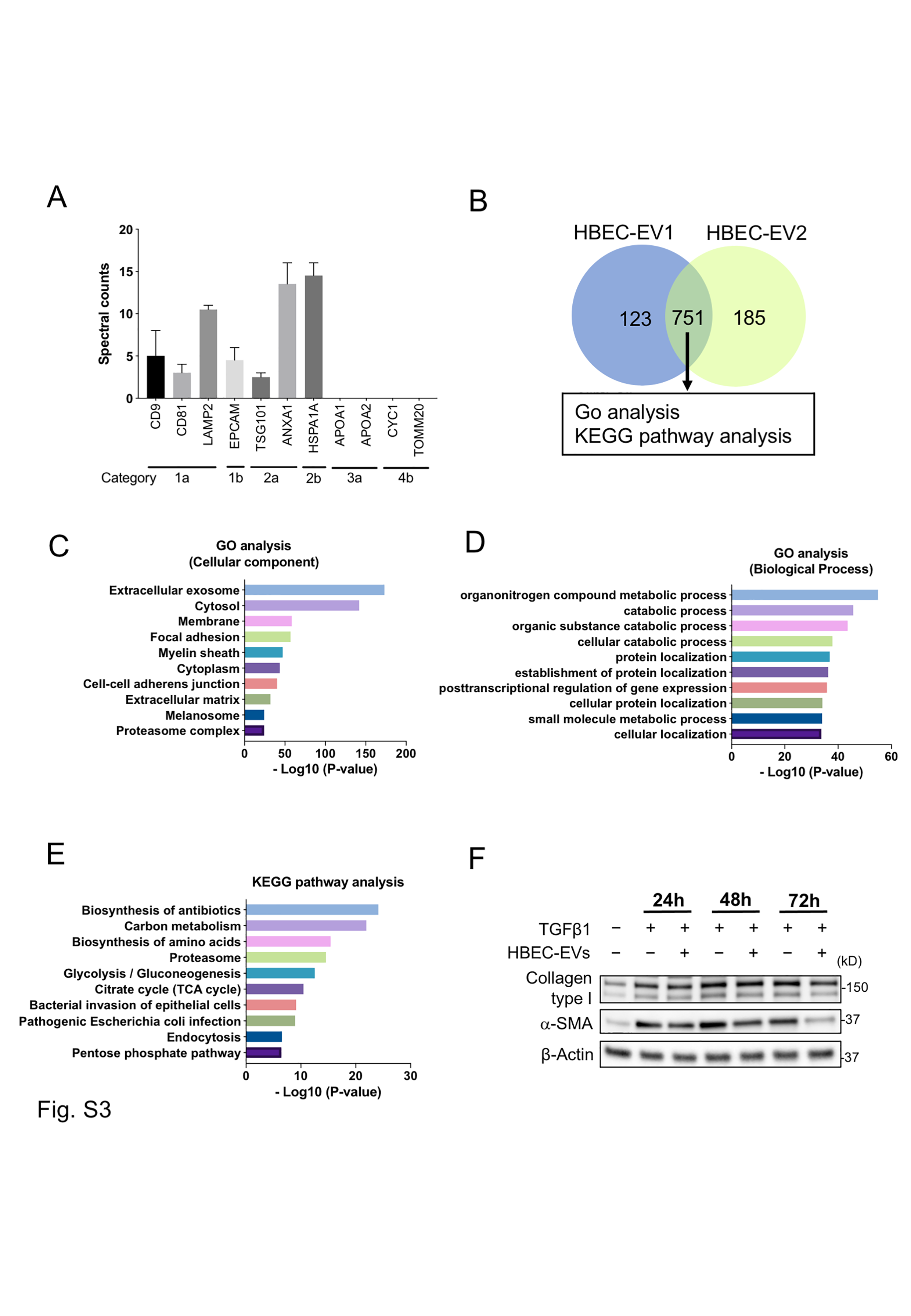

### Fig. S4

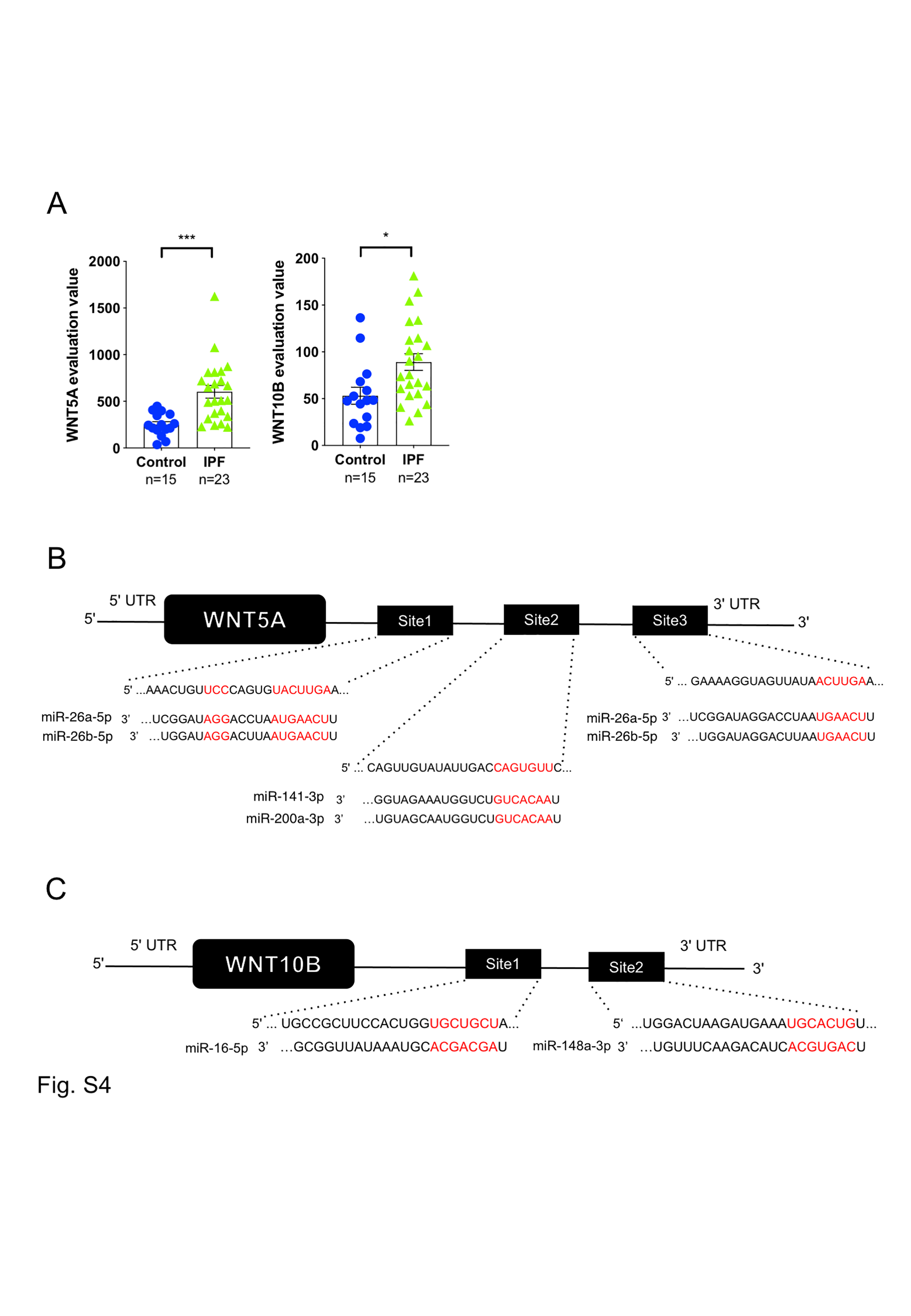

### Fig. S5

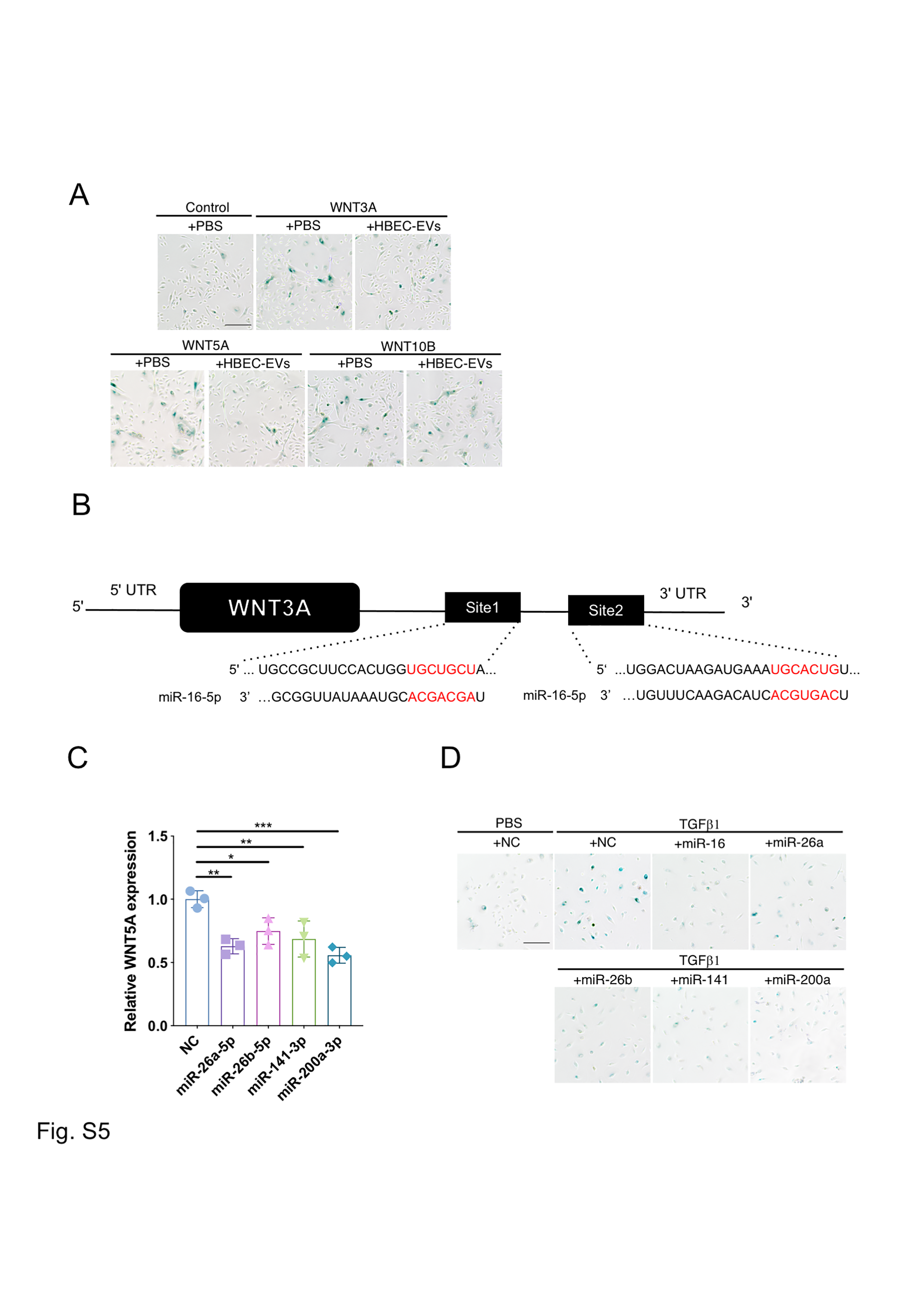

### Fig. S6

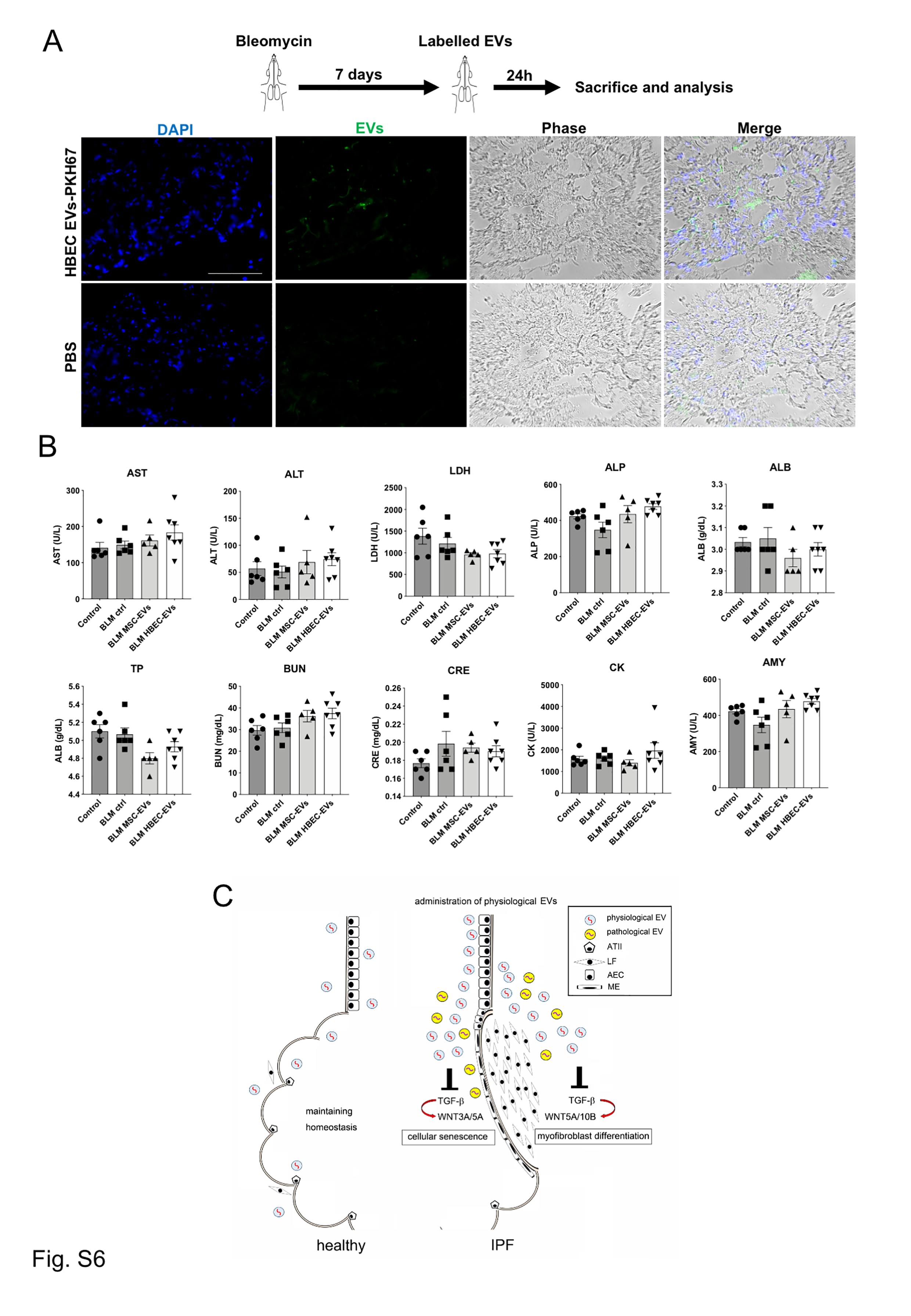

### Table S3

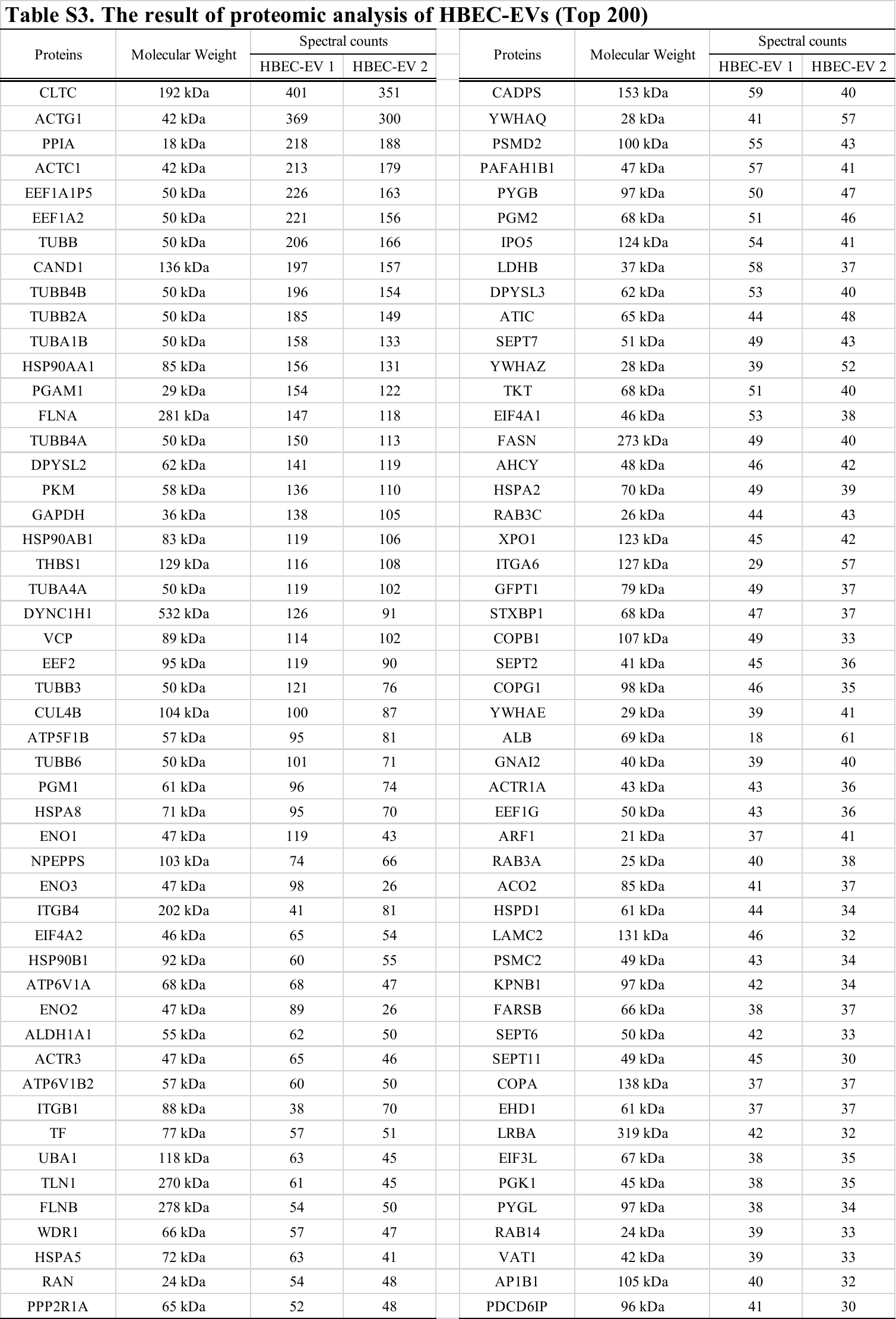

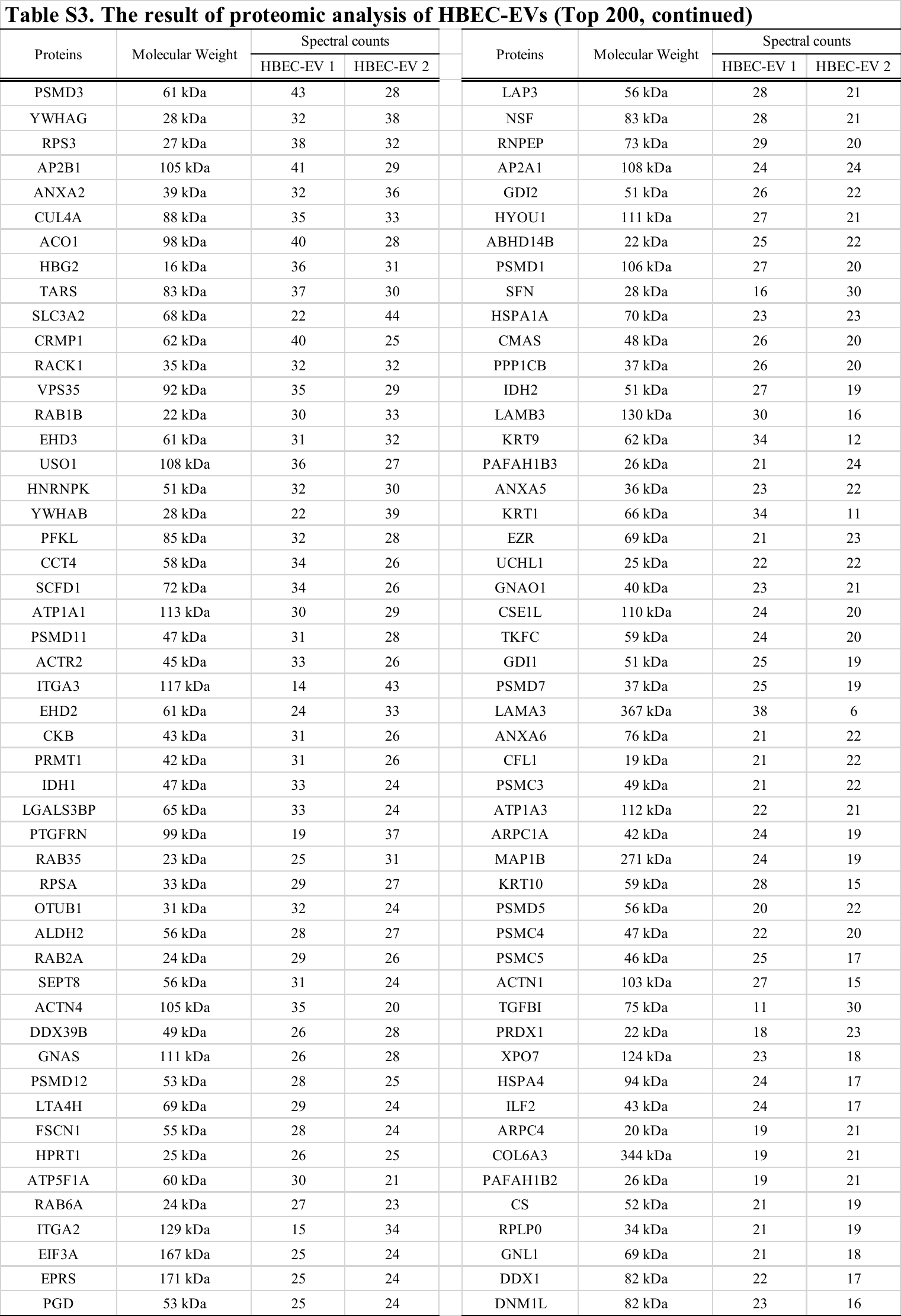
