## Supplementary material for "Human bronchial epithelial cell-derived extracellular vesicle therapy for pulmonary fibrosis via inhibition of TGF-β-WNT crosstalk": Table S1

| **Table S1. Donor characteristics (for lung epithelial cells)** | | |
| --- | --- | --- |
|  | HBECs (n=16) | HSAECs (n=4) |
| Age, years | 67.5 ± 6.9 | 68.8 ± 7.5 |
| Males, % of group | 75.0 | 75.0 |
| SI, pack year | 38.8 ± 35.1 | 13.8 ± 14.7 |
| %VC | 103.5 ± 15.4 | 100.1 ± 17.0 |
| FEV1.0/FVC (%) | 77.8 ± 15.1 | 86.3 ± 20.2 |
| Abbreviations: HBEC, human bronchial epithelial cell; HSAEC, human small airway epithelial cell; SI, smoking index; VC, vital capacity; FEV1.0, forced expiratory volume in 1 second; FVC, forced vital capacity. | | |
