## Supplementary material for "Human bronchial epithelial cell-derived extracellular vesicle therapy for pulmonary fibrosis via inhibition of TGF-β-WNT crosstalk": Table S2

| **Table S2. Donor information for lung fibroblasts from patients with or without IPF.** | | | | | | | | | | |
| --- | --- | --- | --- | --- | --- | --- | --- | --- | --- | --- |
|  | IPF #1 | IPF #2 | IPF #3 | IPF #4 | IPF #5 | Non-IPF #1 | Non-IPF #2 | Non-IPF #3 | Non-IPF #4 | Non-IPF #5 |
| Age | 68 | 74 | 82 | 78 | 72 | 72 | 76 | 56 | 61 | 72 |
| Sex | Male | Male | Male | Male | Male | Male | Male | Male | Male | Male |
| SI, pack year | 300 | 0 | 75 | 1160 | 1530 | 0 | 38 | 35 | 0 | 35 |
| %VC | 68.9 | 64 | 63.4 | 106.5 | 68.2 | 103.2 | 121.6 | 120.2 | 121.5 | 102.4 |
| FEV1.0/FVC (%) | 75.9 | 73.5 | 96.7 | 77.85 | 84.6 | 103.4 | 78.2 | 109.0 | 75.0 | 119.7 |
| Abbreviations: IPF, idiopathic pulmonary fibrosis; SI, smoking index; VC, vital capacity; FEV1.0, forced expiratory volume in 1 second; FVC, forced vital capacity. | | | | | | | | | | |
