## Supplementary material for "Human bronchial epithelial cell-derived extracellular vesicle therapy for pulmonary fibrosis via inhibition of TGF-β-WNT crosstalk": Table S4

| **Table S4. qRT-PCR primers for mRNA** | | | |
| --- | --- | --- | --- |
| Gene symbol | Name | Assay ID | Company |
| ACTB | actin beta | Hs01060665_g1 | Thermo Fisher |
| WNT1 | Wnt family member 1 | Hs00180529_m1 | Thermo Fisher |
| WNT3A | Wnt family member 3A | Hs00263977_m1 | Thermo Fisher |
| WNT5A | Wnt family member 5A | Hs00998537_m1 | Thermo Fisher |
| WNT10B | Wnt family member 10B | Hs00928823_m1 | Thermo Fisher |
