## Supplementary material for "Human bronchial epithelial cell-derived extracellular vesicle therapy for pulmonary fibrosis via inhibition of TGF-β-WNT crosstalk": Table S5

| **Table S5. qRT-PCR primers for miRNA** | | |
| --- | --- | --- |
| Name | Assay ID | Company |
| hsa-miR-16-5p | 000391 | Thermo Fisher |
| hsa-miR-26a-5p | 000405 | Thermo Fisher |
| hsa-miR-26b-5p | 000407 | Thermo Fisher |
| hsa-miR-141-3p | 000463 | Thermo Fisher |
| hsa-miR-148a-3p | 000470 | Thermo Fisher |
| hsa-miR-200a-3p | 000502 | Thermo Fisher |
| RNU6B | 001093 | Thermo Fisher |
