## Supplementary Information for "Human bronchial epithelial cell-derived extracellular vesicle therapy for pulmonary fibrosis via inhibition of TGF-β-WNT crosstalk"

**Table of Contents**

Fig. S1. The characterization of lung epithelial cell-derived EVs

Fig. S2. Investigation of signaling pathways targeted by HBEC EVs

Fig. S3. Proteomic analysis of HBEC EVs

Fig. S4. Binding sites of candidate miRNAs in the 3’UTR of WNT5A and WNT10B mRNA

Fig. S5. Binding sites of miR-16-5p in the 3’UTR of WNT3A mRNA and representative images of SA-β-gal staining

Fig. S6. The uptake of HBEC EVs into lung cells *in vivo* and blood biochemistry analysis

Table S1. Donor characteristics (for lung epithelial cells)

Table S2. Donor information for lung fibroblasts from patients with or without IPF

Table S3. The result of proteomic analysis of HBEC EVs (Top 200)

Table S4. qRT-PCR primers for mRNA

Table S5. qRT-PCR primers for miRNA

**Figure S1. The characterization of lung epithelial cell-derived EVs.**

**A** Nanoparticle tracking analysis showing the particle size of HBEC EVs, BEAS-2B EVs and HSAEC EVs. The vertical axis in the graphs shows the number of EV particles (x 10^6^) / mL, and the horizontal axis indicates the particle size (nm) of EVs.

**B** Representative immunoblot of conventional EV markers for whole cell lysates and EVs of HBECs, BEAS-2B cells and HSAECs. 1 μg/lane.

**C** Comparison of protein quantity in HBEC EVs, BEAS-2B EVs and HSAEC EVs. The vertical axis in the graphs shows the amount of EV protein / cell. Error bars represent SEM. ***P*<0.005, ****P*<0.0005. NS; not significant.

**D** Comparison of particle numbers for HBEC EVs, BEAS-2B EVs and HSAEC EVs. The vertical axis in the graphs shows the number of EV particles / cell. Error bars represent SEM. **P*<0.05, ****P*<0.001. NS; not significant.

**E** Comparison of protein concentration per particle for HBEC EVs, BEAS-2B EVs and HSAEC EVs. The vertical axis in the graphs shows the amount of EV protein / cell. Error bars represent SEM. NS; not significant.

**F** Quantification of type I collagen /β-actin and α-SMA /β-actin in LFs treated for 24 h with PBS, HBEC EVs, HSAEC EVs, BM-MSC EVs, LF EVs, HUVEC EVs, THP1 EVs, A549 EVs, PC9 EVs, or PC14 EVs in the presence or absence of TGF-β1 (2 ng/ml). EVs were added to the medium at a concentration of 2×10^9^ particles per ml. Protein samples were collected 72 h after stimulation. *p<0.05, **p<0.005, ***p<0.001.

**G** Quantification of type I collagen, α-SMA, and β-actin in LF treated for 24 h with PFD (10 or 500 μg/ml) or nintedanib (NTD) (100 or 1000 nM) in the presence or absence of TGF-β1 (2 ng/ml). Protein samples were collected 72 h after stimulation. *p<0.05, **p<0.005, ***p<0.001.

**H** Quantification of p21/β-actin expression in HBECs incubated for 48 h with PBS, HBEC EVs, HSAEC EVs and BM-MSC EVs in the presence or absence of TGF-β1 (2 ng/ml) for 48 h. EVs were added to the medium at a concentration of 2×10^9^ particles per ml. Protein samples were collected 96 h after stimulation. **p<0.005, ***p<0.001.

**Figure S2. Investigation of signaling pathways targeted by HBEC EVs.**

**A** Left and upper right: Representative immunoblot and quantitative analysis showing the amount of phospho-SMAD2, SMAD2, phospho-SMAD3, SMAD3, phospho-AKT, AKT, phospho-p38MAPK, p38MAPK, phospho-ERK1/2, ERK1/2 and β-actin in LFs treated for 60 minutes with PBS or HBEC-EVs (10 μg/ml) in the presence or absence of TGF-β1 (2 ng/ml). Lower right: Representative immunoblot and quantitative analysis showing the amount of NOX4 and β-actin in LFs treated for 24 h with PBS or HBEC-EVs (10 μg/ml) in the presence or absence of TGF-β1 (2 ng/ml). *p<0.05, **p<0.005, ***p<0.001.

**B** Representative immunoblot showing the amount of p21 and β-actin in HBECs incubated for 48 h with PBS or BEAS-2B EVs (10 μg/ml) in the presence or absence of TGF-β1 (2 ng/ml). Protein samples were collected 48 h after stimulation.

**C** Heat map of RNA-seq analysis showing differences in expression of the fibrogenic components in LFs treated for 48 h with PBS, HBEC EVs (10 μg/ml) or BEAS-2B EVs (10 μg/ml).

**D** Gene Set Enrichment Analysis (GSEA) of HBECs treated for 48 h with PBS or HBEC EVs (10 μg/ml), revealing downregulation of TGF-β1, extracellular matrix, extracellular matrix structural constituent and collagen fibril organization. NES: a normalized enrichment score. The p-value was calculated by GSEA.

**E** GSEA of HBECs treated for 48 h with PBS and BEAS-2B EVs (10 μg/ml), revealing downregulation of TGF-β1, extracellular matrix, extracellular matrix structural constituent and collagen fibril organization. NES: a normalized enrichment score. The p-value was calculated by GSEA.

**Figure S3. Proteomic analysis of HBEC EVs.**

**A** The spectral counts of HBEC EV protein composition. Typical proteins were selected to demonstrate the presence of EVs according to a MISEV 2018 position statement(1).

**B** Venn diagrams showing the overlap between proteins from two different HBEC EVs preparations.

**C, D** GO analysis of proteins co-expressed in two different HBEC EV preparations.

**E** Kyoto Encyclopedia of Genes and Genomes (KEGG) pathway analysis of proteins co-expressed in two different HBEC EV preparations.

**F** Time course response of HBEC EV effect on TGF-β1 stimulation. Representative immunoblot showing the amount of type I collagen, α-SMA, and β-actin in LFs treated with PBS or HBEC EVs in the presence or absence of TGF-β1 (2 ng/ml).

**Figure S4. Binding sites of candidate miRNAs in the 3’UTR of *WNT5A* and *WNT10B* mRNA.**

**A** WNT5A and WNT10B evaluation value in the mRNA microarray dataset GSE10667 from the Gene Expression Omnibus (GEO) database. *p<0.05, **p<0.005, ***p<0.001.

**B** A schematic representation of binding sites for miR-26a-5p, miR-26b-5p, miR-141-3p and miR-200a-3p in the 3’UTR of *WNT5A* mRNA.

**C** A schematic representation of binding sites for miR-16-5p and miR-148a-3p in the 3’UTR of *WNT10B* mRNA.

**Figure S5. Binding sites of miR-16-5p in the 3’UTR of *WNT3A* mRNA and representative images of SA-β-gal staining.**

**A** Representative images of SA-β-gal staining in HBECs treated for 48h with PBS or HBEC EVs (10 mg/ml) in the presence or absence of WNT3A (200 pg/ml), WNT5A (400 pg/ml), or WNT10B (200 pg/ml). Staining was performed 96 h after stimulation. Scale bars, 200 μm.

**B** A schematic representation of binding sites for miR-16-5p in the 3’UTR of *WNT3A* mRNA.

**C** qRT-PCR analysis of *WNT5A* mRNAs in HBECs transfected with candidate target miRNA mimics or negative control. **P*<0.05, ***P*<0.005, ****P*<0.001.

**D** Representative images of SA-β-gal staining in HBECs treated with validated miRNA mimics or negative control in the presence or absence of TGF-β1 (2 ng/ml). Staining was performed 96 h after stimulation. Scale bars, 200 μm.

**Figure S6. The uptake of HBEC EVs into lung cells *in vivo* and blood biochemistry analysis.**

**A** Representative microscopic images of sections of lung treated with PKH67-labelled HBEC EVs (DAPI: nuclei, green: EVs). HBEC EVs were labelled with PKH67 and 2×10^9^ particles were injected intratracheally. Scale bars, 100 μm.

**B** Evaluation of systemic toxicity of intratracheal administration of HBEC EVs or BM-MSC EVs in mice. Blood levels were determined for aspartate transaminase (AST), alanine transaminase (ALT), lactate dehydrogenase (LDH), alkaline phosphatase (ALP), albumin (ALB), total protein (TP), blood urea nitrogen (BUN), creatinine (CRE), creatine kinase (CK) and amylase (AMY). Each data point represents data from one animal; control n=6, bleomycin (BLM) n=6, BM-MSC EVs n=5, HBEC EVs n=7.

**C** Schematic diagram of the proposed mechanisms for fibrosis resolution by HBEC EVs. HBEC EVs can be transferred to both epithelial cells and fibroblasts in alveolar regions as a means of maintaining microenvironment homeostasis in healthy lungs. On the other hand, EVs derived from metaplastic epithelial cells (ME) present during bronchiolization may have a pathological profibrotic role during the aberrant wound healing process in IPF. This can be controlled by administration of a sufficient quantity of physiological HBEC EVs as a means of regulating TGF-β1 and WNT signaling pathways.
